# Action observation reveals a network with divergent temporal and parietal lobe engagement in dogs compared to humans

**DOI:** 10.1101/2023.10.02.560112

**Authors:** Magdalena Boch, Sabrina Karl, Isabella C. Wagner, Lukas L. Lengersdorff, Ludwig Huber, Claus Lamm

**Affiliations:** Social, Cognitive and Affective Neuroscience Unit, Department of Cognition, Emotion, and Methods in Psychology, Faculty of Psychology, University of Vienna, 1010 Vienna, Austria; Department of Cognitive Biology, Faculty of Life Sciences, University of Vienna, 1090, Vienna, Austria; Comparative Cognition, Messerli Research Institute, University of Veterinary Medicine Vienna, Medical University of Vienna and University of Vienna, 1210 Vienna, Austria

**Author notes:** These authors share senior authorship (Ludwig Huber, Claus Lamm).

**Keywords:** comparative neuroimaging, action observation network, social cognition, dogs, humans

## Abstract

Action observation is a fundamental pillar of social cognition. Neuroimaging research has revealed a human and primate action observation network (AON) encompassing fronto-temporo-parietal areas with links to a species’ imitation tendencies and relative lobe expansion. Dogs (*Canis familiaris)* have good action perception and imitation skills and a less expanded parietal than temporal lobe, but their AON remains unexplored. We conducted a functional MRI study with 28 dogs and 40 humans and found functionally analogous involvement of somatosensory and temporal brain areas of both species’ AONs and responses to transitive and intransitive action observation in line with their imitative skills. However, activation and task-based functional connectivity measures suggested significantly less parietal lobe involvement in dogs than in humans. These findings advance our understanding of the neural bases of action understanding and the convergent evolution of social cognition, with analogies and differences resulting from similar social environments and divergent brain expansion, respectively.

## 1 Introduction

Other individuals’ actions contain a wealth of social information. For example, we may use them to infer others’ intentions^1^ or learn a new skill by imitating their actions^2^. Perceiving others’ actions also plays an integral part in social interactions, such as cooperation^3^ and therefore helps foster long-term social relationships.

Neuroimaging research in humans has consistently revealed that the observation of transitive (i.e., goal-directed) and intransitive (i.e., goal-absent) actions is underpinned by the action observation network (AON; see e.g.^4,5^ for meta-analysis or review). This distributed network comprises the inferior occipito-temporal ventral visual pathway housing face-and body-sensitive areas and a lateral occipito-temporal pathway associated with the perception of dynamic aspects of social cues and action features (see ^6–8^ for reviews). The parietal dorsal visual pathway, including especially the inferior parietal lobule, is also part of the AON and associated with “vision-to-action” brain functions that guide reach-to-grasp actions (see e.g.,^9^ or ^8^ for review). Finally, the AON involves sensory-motor areas, such as the premotor cortex, which, as the parietal regions, are also active during the observation and execution of the same action (i.e., so-called "mirror neurons" ^5,10,11^).

A similar network, such as the AON, is also present in non-human primates. This network, however, shows species differences that provide important insights into the evolution of the human AON. In rhesus macaques (*Macaca mulatta*) and chimpanzees (*Pan troglodytes*), action observation elicits the strongest activation in frontoparietal regions^12,13^, but common marmosets (*Callithrix jacchus*) have stronger frontotemporal activation^14^. In Old World monkeys, apes and humans, the frontal and parietal lobes expanded disproportionally compared to the other lobes. In contrast, other primate (including marmosets) and mammalian species have more expanded occipital and temporal lobes^15^. Thus, relative lobe expansions might have led to differences in activation patterns across these primate species.

The distributed activation pattern in the human AON has been linked to more pronounced connections (i.e., white matter pathways) between the parietal and especially the inferior temporal cortex, compared to chimpanzees and macaques with the least pronounced connections^16^. In humans and chimpanzees, but not in macaques, the AON is similarly active during both transitive and intransitive action observation^4,12^ (or see^17^ for review). The differences in structural connectivity and activation for transitive and intransitive actions have been associated with the species’ ability to learn tool-using behaviours through social learning and their tendency to imitate actions^16^. Humans and (less successfully) chimpanzees can acquire tool use through social learning, but macaques cannot as they are emulators focussing on the action outcome^16,18–20^. Recent findings in marmosets^14^, which are not known as tool-users either but display true imitation of behaviours^21,22^, make the suggested relationship between comparable activation during transitive and intransitive action observation and the tendency to imitate actions of others less clear. Here, transitive compared to intransitive action observation led to stronger activation, but the marmosets also showed distributed activation in temporal lobe areas during action observation.

Another approach to studying the human AON’s evolutionary history and its link to differential lobe expansion, imitation and object-manipulating behaviours is to search for convergent evolution in a different animal lineage. So far, research in non-primate species has focused on investigations of mirror neurons in the motor and anterior cingulate cortex of rats^23,24^ and in the forebrain of various songbirds^25–27^. However, due to the nature of the electrophysiological recordings of these studies, they were not able to explore whole-brain networks that underpin action observation and thus remain agnostic to putative further functional analogies with humans or non-human primates. Dogs (*Canis familiaris*) represent a highly relevant additional species for comparative research on action observation for several reasons. First, they do not engage in tool-using behaviours but demonstrate good action perception and imitation skills. For example, dogs can perceive actions as goal-directed^28^, and they can imitate actions of dogs and humans^29–33^, and even over-imitate actions demonstrated by their primary human caregivers^34–36^. Second, dogs have a temporal lobe, which evolved in carnivorans independently of primates^37,38^, but as marmosets, they do not have a significantly expanded parietal lobe^15^. Third, they can be trained to participate awake and unrestrained in functional MRI studies^39^, allowing us to conduct comparative studies on dogs and humans using largely identical scanning conditions. Hence, dogs constitute an excellent model for studying the evolutionary history of the human AON and the relationship to social learning, tool-use behaviours, and differential brain expansion.

In the present comparative functional MRI study, we thus performed the first investigation of the dog action observation network (AON). We aimed to identify functional analogies and divergencies between the dog and human AONs by applying univariate activation and task-based functional connectivity analyses^40,41^. We used experimental manipulations that allowed us to assess AON engagement to intransitive vs. transitive actions and actions performed by con-vs. heterospecifics and controlled for low-level visual aspects.

To identify potential functional analogues between the two species, we first assessed activation in brain areas involved in sensory-motor processes in both species (e.g., dog pre-and postcruciate gyrus and human inferior lateral gyrus; see **Supplementary Figure S1A** for details). Second, we hypothesized that the dog action observation network, as in humans, includes occipito-temporal brain areas associated with face-and body perception (i.e., agent areas), as well as areas involved in processing of dynamic aspects of social cues and action features^6–8^. First evidence suggests that the dog agent areas are housed in the occipito-temporal ectomarginal, the mid and caudal suprasylvian gyrus, but results have been mixed^42–49^. Brain areas associated with the processing of dynamic aspects have yet to be investigated. Thus, to test our hypothesis, we used a functional localizer task to identify the agent areas within the dog AON and disentangle them from areas sensitive to the action features. Third, in light of the species’ imitation and action-matching skills^30,35,50,51^, we predicted comparable activation of the dog and human action observation networks while observing transitive and intransitive actions. Fourth, regarding their close bond, we also explored how the species’ action observation network responded to actions performed by each other vs. conspecifics (i.e., own species).

However, we also expected functional divergencies between the dog and human AON based on prior comparative research with non-human primates^12–14^. Considering the dogs’ and humans’ differential relative lobe expansion^15^, we predicted the strongest engagement of the dog temporal lobe during action observation and less parietal lobe involvement than in humans. We quantitatively tested this hypothesis by first comparing the extent of active voxels in both species’ parietal and temporal lobes during action observation. Then, we also investigated task-based functional connectivity between the primary visual cortex (V1) and each lobe to investigate a potentially greater information exchange with the temporal than parietal lobe in dogs.

## 2 Results

We collected comparative functional MRI data from 28 pet dogs and 40 human participants while viewing videos of dogs and humans performing transitive or intransitive actions, as well as low-level visual and motion controls (see **Figure 1**). We implemented the study in a block design and used a tailored haemodynamic response function^52^ (HRF) for the dog data to improve detection power. All reported statistical tests were corrected for multiple comparisons.

**Figure 1.**
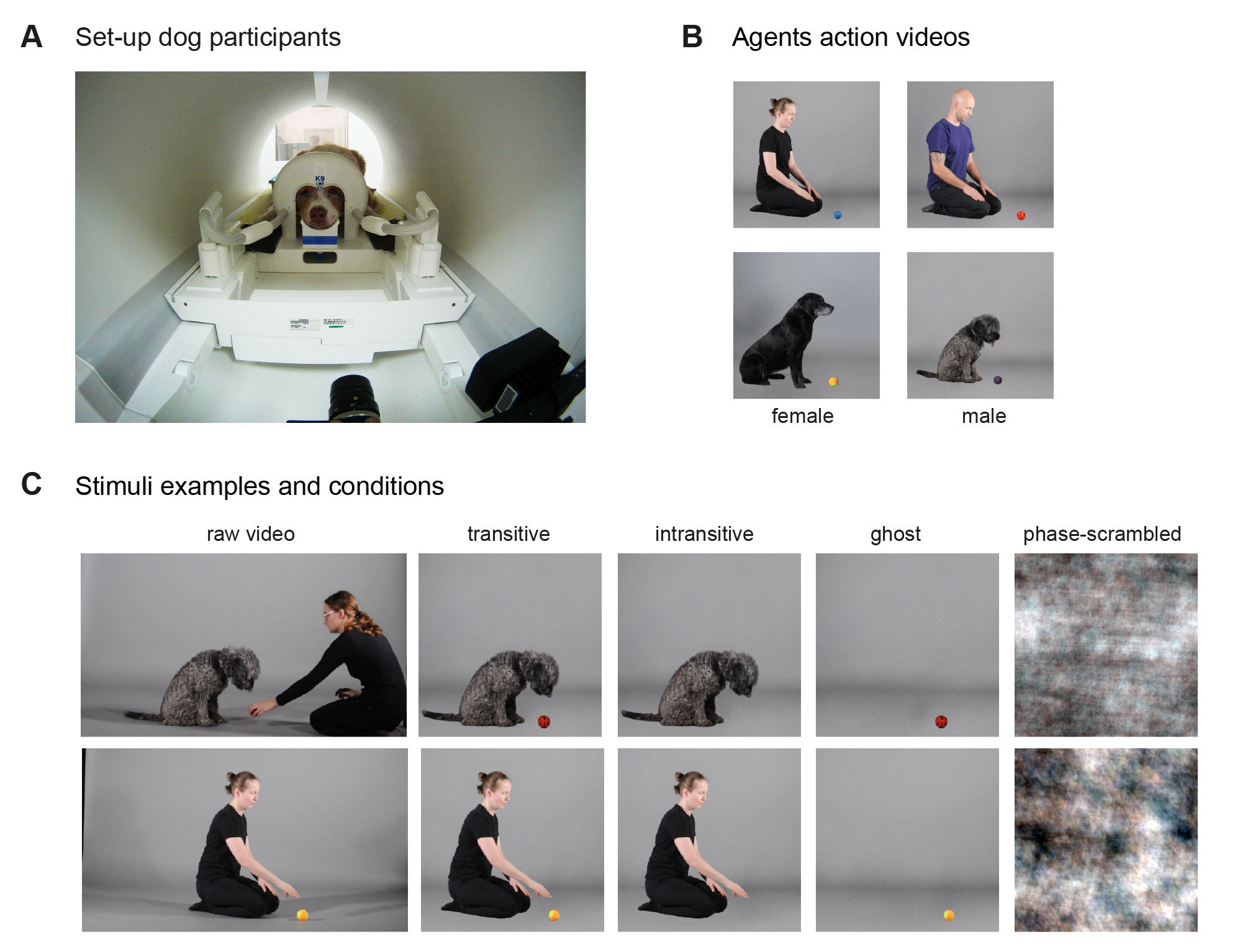
Experimental design. (**A**) Dogs were positioned in sphinx position with their head placed in a custom-made 16-channel (k9) head coil^53^. For human participants, data were obtained with the same Siemens 3T scanner but with a 32-channel human head coil in a supine position and with stimuli presented via a back-projection mirror mounted on the coil (not shown). (**B**) All participants underwent scanning under the same conditions and using the exact same video stimuli depicting four unfamiliar agents (2 humans, 2 dogs) picking up different toy balls (4 videos/agent). (**C**) The experimental design of the study entailed the two main factors, action (transitive, intransitive) and agent (dog, human) and two additional control conditions: ghost and scrambled (transitive) videos. All conditions were derived from the same raw videos by cutting out the dog trainer if applicable (transitive), the toy (intransitive), agent (object motion) or by phase-scrambling the transitive video. Stimuli were presented in a block design (12 s) with four videos per block and data acquisition was split into two 5-min task runs. All example images of the stimulus set are screenshots of material created by the authors, and we have obtained consent from individuals featured in the videos for their public dissemination. Due to bioRxiv’s policy, we had to remove this figure from the manuscript but it is publicly available at the study’s OSF data repository: https://osf.io/z479k/files/osfstorage/651adfdeb0a33e07995921b6

### 2.1 Functionally convergent temporal and somatosensory components of the dog and human action observation network

First, to localize the dog and human action observation network, we investigated activation in response to the observation of transitive (i.e., goal-directed) and intransitive (i.e., goal absent) actions of both species (i.e., average activation of all action conditions) compared to the implicit visual baseline and the object and scrambled motion controls.

Action observation, compared to the implicit visual baseline, led to activation in occipito-temporal and somatosensory cortex in dogs and humans alike (see **Figure 2A**, **Supplementary Tables S1-S2**). In dogs, occipito-temporal clusters expanded across primary (i.e., marginal gyrus) and extrastriate (i.e., splenial, ectomarginal gyrus) visual cortices, as well as higher-order association cortices, including the mid and caudal suprasylvian and the sylvian and ventral caudal composite gyrus (see **Supplementary Figure S2A** for overview of the main gyri and sulci of the dog brain). We also found activation in the secondary somatosensory cortex (i.e., rostral ectosylvian gyrus) but no evidence for the involvement of premotor cortices (i.e., precruciate gyrus). Due to anticipated signal loss in this area caused by dogs’ large nasal cavities (**Supplementary Figure S1B-C**), we also conducted region-of-interest analyses (see below). In humans, occipito-temporal activation also comprised the striate and extrastriate cortices (e.g., MT: middle temporal area / V5, lateral occipital cortex), as well as the inferior temporal cortex (i.e., fusiform gyrus) and posterior superior temporal sulcus (pSTS), and we found activation in human somatosensory cortices (i.e., postcentral gyrus). Exclusively in humans, we observed activation in parietal cortices (i.e., superior and inferior lobules), sub-cortical areas such as the amygdala and thalamus, and (pre-) motor cortices (i.e., inferior frontal and precentral gyrus).

**Figure 2.**
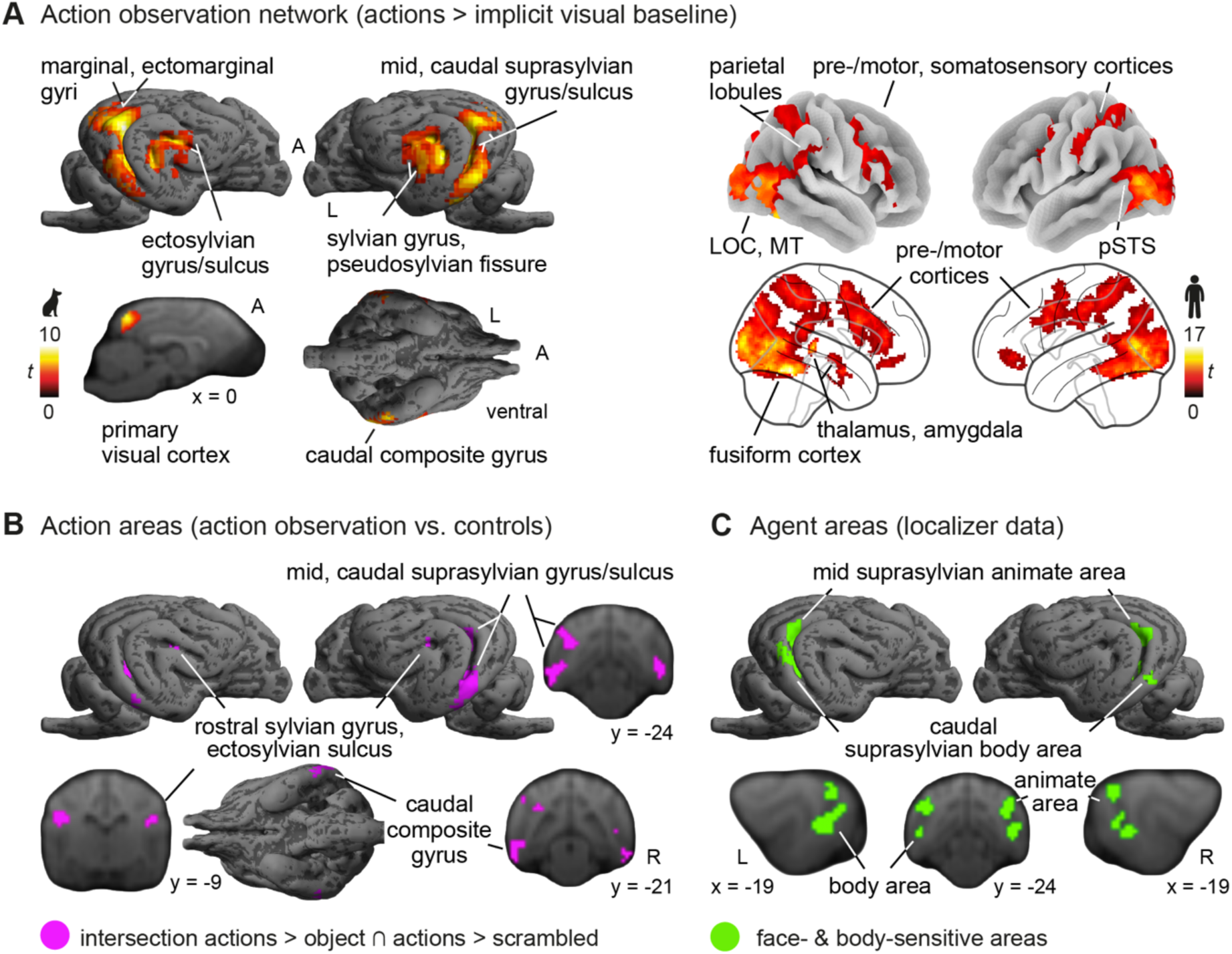
Dog and human action observation networks. (**A**) Action observation (i.e., pooled activation for transitive and intransitive actions of dogs and humans) compared to implicit visual baseline led to functionally analogous activation in dog and human occipito-temporal and somatosensory cortices, but parietal and (pre-)motor activation was only observed in humans (see also **Supplementary Tables S1-S2**). (**B**) Action observation compared to object and scrambled motion (i.e., purple-coloured clusters represent the intersection of both contrasts; see **Supplementary Figure S3** for separate contrasts) revealed activation in the dog caudal composite and rostral sylvian gyrus including the ectosylvian sulcus (henceforth referred to as rostral sylvian area), and the mid and caudal suprasylvian gyrus and sulcus. (**C**) The latter two areas largely overlap with the mid suprasylvian animate area, which is sensitive to faces and bodies compared to inanimate objects and scrambled controls, and the caudal suprasylvian body area, which responds significantly stronger to bodies compared to faces and visual controls as identified with the agent localizer task. Green-coloured clusters represent the animate and body areas (see **Supplementary Figure S8** for details). Results are *p* < .05 FWE-corrected at cluster-level using a cluster-defining threshold of *p* < .005/.001 for dogs/humans. Anatomical nomenclature for all figures refers to the dog brain atlas from Czeibert et al. (2019) normalized to the stereotaxic breed-averaged template space ^55^ and the Harvard-Oxford human brain atlas ^56^. MT, middle temporal visual area (V5); A, anterior; R, right; L, left; *t*, *t*-values. The dog and human icons in A were purchased from thenounproject.com (royalty-free license).

In dogs, activation in the higher-order occipito-temporal association cortices remained after controlling for low-level visual stimulation (i.e., scrambled motion) or object motion (see **Figure 2B** for the intersection of both contrasts), while humans showed activation in temporal, parietal and somatosensory areas. Human premotor activation was only found when contrasting with the scrambled motion control, but not the object motion control (see **Supplementary Figure S3** for each contrast separately).

#### 2.1.1 Functionally analogous temporal areas sensitive to face-and body perception and agents-in-action

Temporal lobe activation in the human action observation network has been categorized as pertaining to two distinct components: face-and body-sensitive areas in the ventral visual pathway and areas sensitive to dynamic visual aspects of social cues and action features in the lateral temporal or third visual pathway (see e.g.,^6–8^ for reviews). To identify the functional analogues of the human face-and body-sensitive areas and disentangle them from potential functional analogues of human lateral temporal brain areas sensitive to action features in dogs, we used data from a functional localizer. This identified an area in the mid suprasylvian gyrus of dogs that was sensitive to faces and bodies and an additional area associated with body perception in the dog caudal suprasylvian gyrus (see **Figure 2C** and **Supplementary Note 1**). These areas overlapped with the mid and suprasylvian clusters in the dog AON (see **Figure 2B**). However, action observation also led to activation in further temporal areas, such as the ventral caudal composite gyrus and a cluster including the rostral sylvian gyrus and ectosylvian sulcus, henceforth referred to as the rostral sylvian area.

#### 2.1.2 No differences in activation for transitive vs. intransitive actions in dogs, and greater activation for dog compared to human actions in both species

Next, we tested our hypothesis of comparable engagement of the dog and human action observation network during transitive and intransitive action observation due to the species’ imitation skills. We further explored how they perceived actions performed by the other species vs. by conspecifics. To address these research questions, we computed a within-subjects full factorial model with the factors *action* (transitive, intransitive) and *agent* (dog, human). Results revealed significant main effects of action in humans, and of agent in both species, but no significant action x agent interaction in either species (see **Figure 3A** and **Supplementary Table S7**).

**Figure 3.**
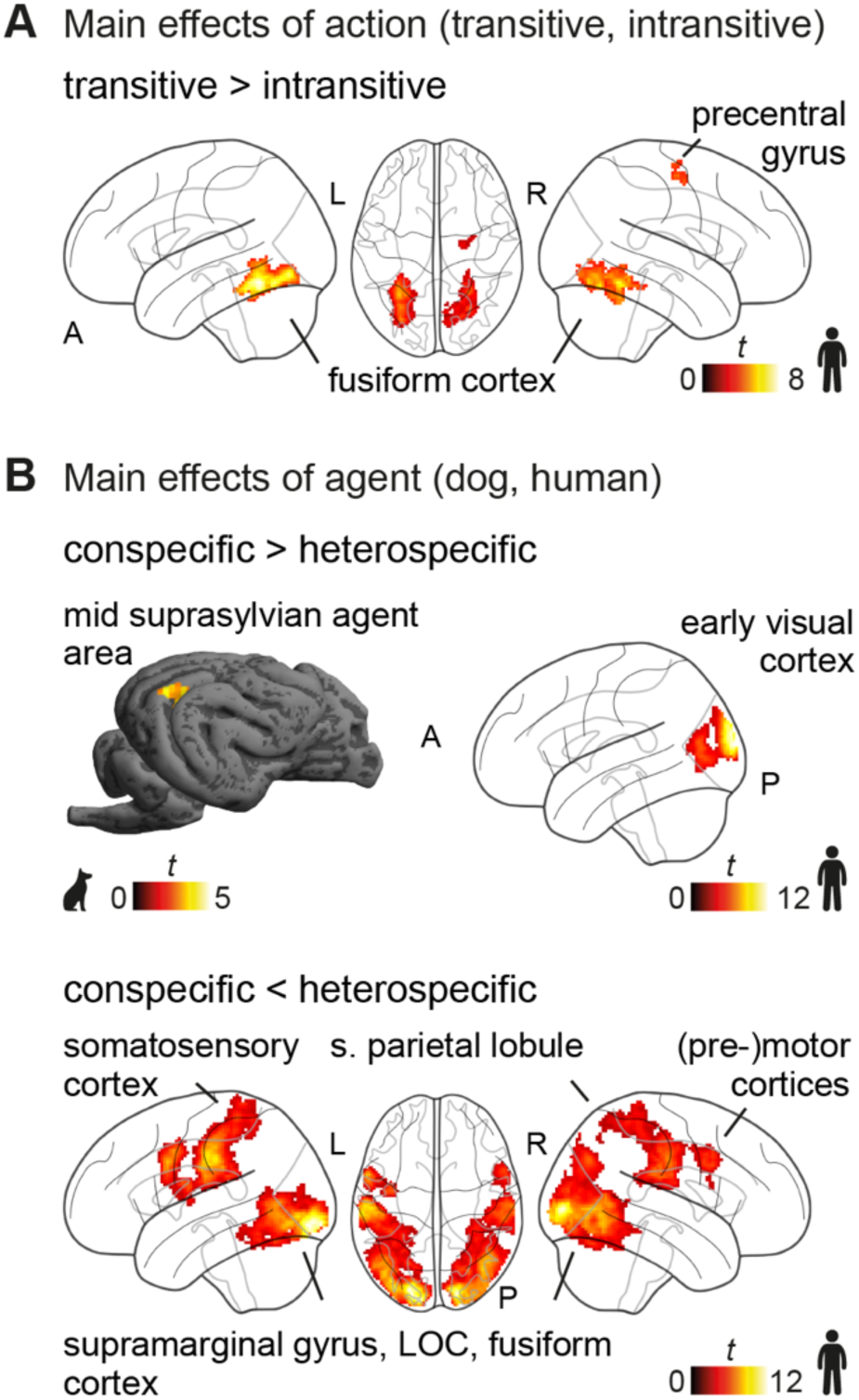
No differences in activation for transitive vs. intransitive actions in dogs and greater activation for observing dog compared to human actions in both species. (**A**) Transitive compared to intransitive actions led to greater activation in the human inferior temporal cortex. (**B**) Observing conspecific compared to heterospecific actions led to greater activation in the dog mid suprasylvian agent area and human early visual cortex. The reversed contrast revealed greater activation in frontoparietal and occipito-temporal cortices in humans but no significant cluster in dogs. Results are *p* < .05 FWE-corrected at cluster-level using a cluster-defining threshold of *p* < .005/.001 for dogs/humans, see **Supplementary Table S7**). LOC, lateral occipital cortex; A, anterior; P, posterior; L, left; R, right; s., superior, *t*, *t*-value. The dog and human icons were purchased from thenounproject.com (royalty-free license).

In more detail, transitive compared to intransitive action observation led to greater activation in the primary somatosensory cortex (i.e., precentral gyrus) and the inferior temporal cortex in humans (**Figure 3A**). We did not find significant activation for the reversed contrast. In dogs, we did not find a significant difference in activation between transitive and intransitive actions. Activation maps for each condition (compared to the implicit visual baseline) confirmed similar activation for transitive and intransitive actions in dogs (see **Supplementary Figure S4-S5** for activation of each condition-of-interest > implicit baseline for both species).

Observing conspecific compared to heterospecific actions resulted in greater activation in the left mid suprasylvian animate area of dogs and the primary visual cortex of humans (**Figure 3B**). The reversed contrast did not reveal any significant clusters. In humans, observing heterospecific (i.e., dog) compared to conspecific actions resulted in greater activation in clusters expanding across the frontoparietal (i.e., pre-and postcentral gyrus, bilateral supramarginal gyrus) and occipital-temporal cortices (e.g., MT, fusiform gyrus, inferior occipital gyrus), largely overlapping with the activation observed for action observation in (see e.g., **Figure 2A**).

### 2.2 Region-of-interest analysis reveals somatosensory but no (pre-) motor activation during action observation in dogs

We also conducted a (planned) region-of-interest (ROI) analysis in the dog sensory-motor cortices to test our hypothesis of somatosensory and (pre-)motor engagement during action observation in dogs, functionally analogous to humans.

The linear mixed model (LMM) yielded no significant changes in activation levels for action observation compared to object or scrambled motion in the dog pre-and motor cortices (see **Figure 4A** and **Supplementary Table S5**). Visual exploration of the data revealed that only eight out of the *N =* 28 dogs displayed activation levels higher for actions compared to both visual controls. As expected, the temporal signal-to-noise ratio in dog frontal lobes was decreased due to air-filled cavities (i.e., sinuses) partly affecting the precruciate gyrus (i.e., premotor and secondary motor cortex see **Supplementary Figure S1B-C**). The ROI analysis revealed, however, somatosensory involvement during action observation in dogs. We found greater activation for action observation compared to object motion in the dog primary somatosensory cortex (rostral suprasylvian gyrus) and greater activation for action observation compared to both controls in the dog secondary somatosensory cortex (rostral ectosylvian gyrus). Activation levels in dog sensory-motor cortices were not modulated by agent or action type (see **Supplementary Table S5**).

**Figure 4.**
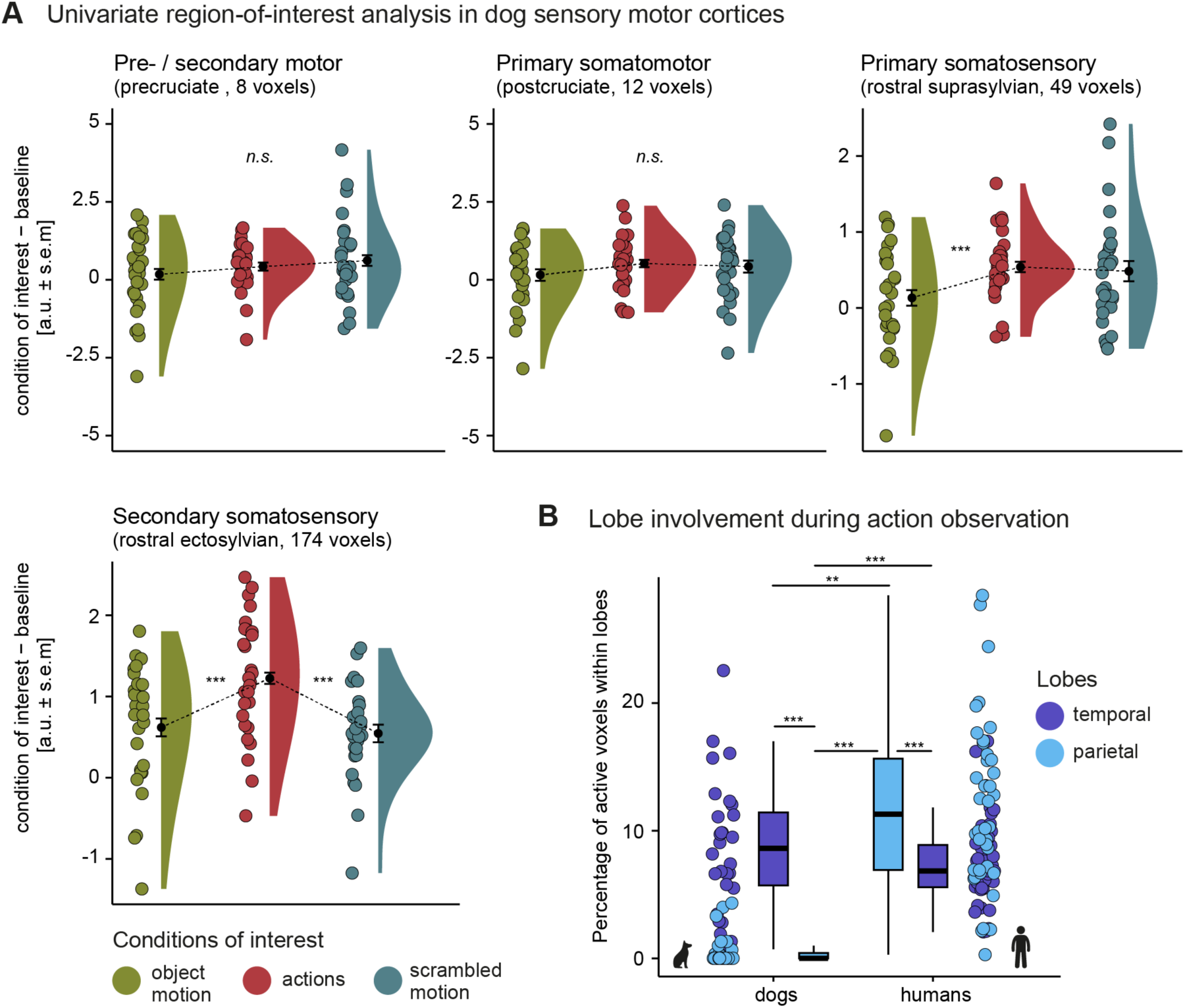
**Somatosensory but no (pre-) motor activation in dogs and significantly more temporal than parietal activation in the dog compared to human action observation network.** (**A**) Action observation compared to object and scrambled motion control did not lead to differences in activation levels in dog (pre-) motor cortices but in the rostral suprasylvian primary somatosensory and rostral ectosylvian secondary somatosensory cortex. The raincloud plots^57^ show the group mean activation measured in arbitrary units (a.u.) with error bars indicating the standard error of the mean (s.e.m.), individual means (coloured dots) and density plots (half violins). (**B**) Action observation resulted in a significantly higher percentage of active voxels in the human compared to the dog parietal lobe. Dogs had significantly more active voxels in the temporal than the parietal lobe, while humans showed a reversed pattern. Discrepancies between lobes were more significant in dogs, with most having 0% active parietal voxels. We determined active voxels for the individual action observation contrast maps (i.e., all conditions displaying an agent acting > implicit baseline) by applying cluster-level inference (*p* < .005/.001 dogs/humans) and a cluster probability of *p* < .05 FWE corrected (see also **Supplementary Table S6**). The boxplots show the median percentage (black horizontal line), interquartile range (box), the lower/upper adjacent values (whiskers), and are accompanied by coloured dots representing the individual percentages. Planned comparisons were false discovery rate (FDR) corrected to control for multiple comparisons. ** *p* < .01, *** *p* < .001; n.s., not significant. The dog and human icons in A were purchased from thenounproject.com (royalty-free license).

### 2.3 Divergent patterns of parietal and temporal engagement in dogs and humans

As the final analysis step, we tested our hypothesis of differential temporal and parietal lobe engagement in the dog and human action observation networks by first comparing the extent of activation in the two lobes within-and between the two species and second the strength of task-based functional connectivity between the primary visual cortex (V1) and the temporal vs. parietal lobe in dogs and humans.

#### 2.3.1 Significantly more temporal than parietal activation in the dog compared to human action observation network

Whole-brain analyses suggested stronger parietal engagement in humans but not in dogs during action observation (see **Figure 2A**). To go beyond this descriptive evaluation, we quantitatively tested the hypothesis of stronger temporal than parietal lobe activation in dogs by directly comparing the activation extent in the parietal lobes of dogs and humans. To this end, we measured the individual percentages of active voxels within each species’ lobe during action observation compared to the implicit visual baseline. Auditory and somatosensory cortices were excluded from the lobe masks (see **Supplementary Figure S2B** and methods section **cross-species comparison of parietal and temporal cortex involvement** for details).

In line with our hypothesis and confirming the descriptive whole-brain findings, the cross-species revealed significantly higher proportions of active voxels in the temporal compared to the parietal lobe of dogs and significantly higher percentages of active voxels in the parietal visual and multisensory cortices of humans as compared to dogs (see **Figure 4B** and **Supplementary Table S6**). The majority of dogs had 0% active voxels in the parietal lobe. We found the reversed pattern with greater parietal than temporal involvement in humans. However, discrepancies between the engagement of the two lobes were less pronounced, showing more distributed involvement of both lobes. Secondary analyses further confirmed that the observed between and within species differences in activation extent also remained using more liberal thresholds to define active voxels (see **Supplementary Figure S6**).

### 2.4 Greater task-based functional connectivity between the primary visual cortex and temporal compared to parietal lobe in dogs

We then compared task-based functional connectivity between V1 and the temporal and parietal lobe during action observation in both species using generalized psychophysiological interaction (gPPI) analyses^40^. This analysis tested whether the more pronounced temporal lobe engagement in dogs was not only translated into greater activation extent but also greater functional connectivity with V1 compared to the parietal lobe within the dog action observation network.

For each species, we extracted task-based functional connectivity with V1 from anatomical masks in the parietal and temporal lobe. We then aggregated the functional connectivity measures across all masks of the same lobe to (1) compare V1 connectivity between lobes (2) and, in response to action, observation compared to the two control conditions by employing an LMM for each species.

The analysis revealed significantly greater connectivity between V1 and the temporal compared to the parietal lobe in dogs. We did not find a significant difference in connectivity in humans between the two lobes (see **Figure 5A** and **Supplementary Tables S8-S9**). In dogs and humans, connectivity between V1 and the temporal lobe was the strongest during action observation compared to both controls. Parietal lobe connectivity patterns diverged in the two species. V1 connectivity was significantly higher in dogs for object than scrambled motion and action observation. In contrast, we did not find differences in connectivity for object motion and action observation in humans, but both significantly differed from scrambled motion.

**Figure 5.**
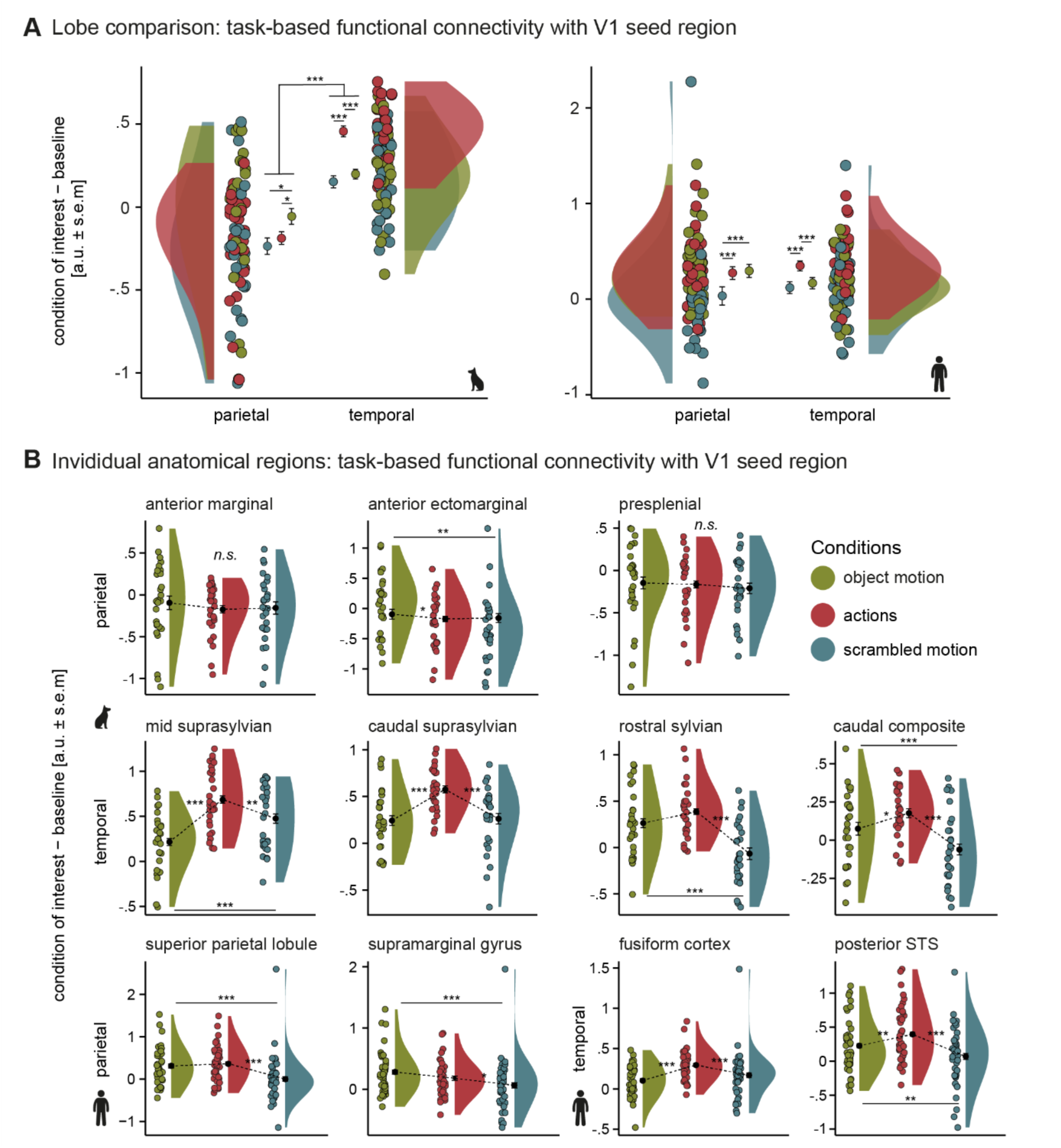
Greater task-based functional connectivity between the primary visual cortex and temporal compared to parietal lobe in dogs. (**A**) Overall, we found greater task-based functional connectivity between the temporal lobe and the primary visual cortex (V1) seed region than the parietal lobe but no differences in V1 connectivity between the two lobes in humans. In dogs and humans, connectivity between V1 and the temporal lobe was significantly greater during action observation compared to both controls. In the parietal lobe, object motion led to higher V1 connectivity compared to scrambled motion and action observation in dogs. In humans, connectivity between V1 and the parietal lobe did not differ between action observation and object motion (see **Supplementary Tables S8-S9**). (**B**) In humans, we found the same connectivity patterns observed on the lobe levels in each individual anatomical mask. In dogs, this was also the case in most anatomical masks, but we did not find a significant effect of condition in the parietal anterior marginal and presplenial gyrus (see **Supplementary Table S10**). V1 task-based functional connectivity during object motion compared to action observation did not significantly differ for the anatomical rostral sylvian mask, but for the more constrained functionally defined rostral sylvian area mask (see **Supplementary Figure S7** for secondary analysis). The raincloud plots^57^ show the group mean task-based functional connectivity with V1 measured in arbitrary units (a.u.) with error bars indicating the standard error of the mean (s.e.m.), individual means (coloured dots) and density plots (half violins). Planned comparisons were false discovery rate (FDR) corrected to control for multiple comparisons. * *p* < .05 ** *p* < .01, ** *p* < .001; n.s., not significant. The dog and human icons in A were purchased from thenounproject.com (royalty-free license).

Next, investigating task-based functional connectivity levels of each anatomical ROI separately, we found the same connectivity patterns as observed on the lobe-level in the dog parietal anterior ectomarginal gyrus and in both human anatomical ROIs (i.e., superior parietal lobule, supramarginal gyrus). We did not find a significant effect of condition in dogs’ parietal anterior marginal and presplenial gyri (see **Figure 5B**, **Supplementary Table S10**). V1 connectivity was the highest during action observation compared to both controls in each dog and human temporal ROI and we found significantly greater connectivity for object compared to scrambled motion in the dog rostral sylvian and caudal composite gyri, and the human posterior STS. In the dog mid suprasylvian gyrus, task-based connectivity was greater for scrambled than object motion. The difference in connectivity between action observation and object motion did not reach significance for the anatomical rostral sylvian gyrus mask.

However, using a more constrained functional mask of the rostral sylvian area derived from the univariate analysis (see **Figure 2B**), we did find a significant difference in connectivity between action observation and both control conditions. All other connectivity results found with the anatomical ROIs in dogs were also replicated using the more constrained functionally defined ROIs (see **Supplementary Figure S7** and **Table S10** for secondary analysis). Finally, results from the localizer task showed that the agent areas also engage in greater information exchange with V1 during face and body compared to the controls, further emphasizing their functionally analogous function of human ventral temporal brain areas (see **Supplementary Note 1** for details).

## 3 Discussion

The major aim of our comparative neuroimaging study was to localize the dog action observation network for the first time and to identify (a) functionally analogous and (b) divergent neural underpinnings with the human action observation network (AON). (a) Our findings revealed multiple functional analogies between the two species, such as the involvement of temporal and somatosensory cortices during action observation compared to the visual control conditions. As expected, in light of the two species’ imitation^33,34,51^, and spontaneous action matching^50^ skills, the dog and human AON responded to the observation of both transitive and intransitive actions with similar activation for both action types and greater activation for transitive actions only in the human inferior temporal cortex. Together with the data from a functional localizer task for the dogs, our results also confirmed our prediction of functionally analogous temporal AON components associated with (1) face and body perception and (2) processing of action features. (b) However, we also identified expected and unexpected divergencies in the species’ AONs. As predicted, based on the differential evolutionary history and relative lobe expansion of the dog and human temporal and parietal lobe^15^, we found more pronounced temporal than parietal lobe involvement in dogs during action observation in terms of the extent of active voxels and task-based functional connectivity with V1. In line with our hypothesis, we found significantly more parietal cortex activation in dogs than in humans during action observation and more distributed involvement of the temporal and parietal lobe within the human AON. Unexpectedly, but likely due to severe signal loss in this area in dogs, we only found premotor activation in response to action observation in humans. Thus, overall, the observed similarities and differences provide novel insights into the evolution of the neural bases of action perception and the link to relative brain expansion and social learning.

Starting with a discussion of the functional analogies, our results show that the AONs of both species include occipital-temporal regions. Observing dogs and humans performing transitive or intransitive actions elicited greater activation and functional connectivity with V1 in the temporal body and agent-sensitive areas of both species (i.e., dog mid and caudal suprasylvian gyrus^46,49^ and human inferior temporal cortex^58^). As expected, action observation also engaged other temporal brain areas sensitive to the dynamic aspects of the visual social cues. In humans, this included the posterior superior temporal sulcus (pSTS), a multi-sensory association cortex involved in action and social perception^59^. In dogs, the present study was the first systematic exploration of these neural bases, and we localized two regions in the temporal lobe: the caudal composite gyrus and an area in the rostral sylvian gyrus, including the rostral ectosylvian sulcus. As the human pSTS, both areas house multisensory cortices and are together with auditory regions (i.e., ectosylvian gyrus) part of a pathway expanding to sensory-motor and prefrontal cortices^60,61^. The sylvian, ectosylvian and suprasylvian gyrus, and premotor and prefrontal cortices (i.e., precruciate, prorean gyrus) have also been identified as a network systematically covarying in size (i.e., grey matter volume) across breeds that correlated with numerous behavioural specializations requiring action perception of other individuals, such as herding, sport fighting or bird flushing and retrieving^62^. Finally, the rostral sylvian gyrus and ectosylvian sulcus are also associated with social vs. non-social interaction^63^ and emotion perception^48^ and these areas showed relative cortical expansion in foxes selected for tameness compared to aggression^64^. This is suggestive of the rostral sylvian area of the dog AON playing a crucial role in multi-sensory social information integration functionally analogous to the human lateral temporal visual pathway^6,7^.

In line with our prediction based on the species’ imitation skills, we further observed that the dog and human AONs were similarly engaged in observing transitive and intransitive actions. We did not find a significant difference between the observation of transitive vs. intransitive action observation in dogs, and no pronounced differences in humans, with greater activation for transitive actions only in parts of the fusiform cortex. Since large parts of the species’ AONs evolved independently in primates and carnivorans^37,38^, our functionally analogous findings are likely the product of convergent evolution. Research suggests that dogs have inherited the ability to copy the behaviours of others already from their wild ancestors, as their closest non-domesticated relative, the grey wolf (*Canis lupus*), possesses complex social abilities^65^ (and see e.g. ^66,67^ for review). Wolves live in family units with strong social bonds and hunt, rear their offspring, and protect their pack together^68,69^. Close cooperation, which often requires accurately and dynamically predicting others’ actions, is thus critical for survival^70^. Therefore, the dog AON might have already evolved in their close, non-domesticated ancestors, but as a result of domestication is now predominantly relevant for cooperating with their human caregivers.

Our findings also revealed important divergencies in the neural bases of action observation, which are of particularly conceptual relevance. Based on prior research with non-human primates^12–14^, we had predicted a notable discrepancy in parietal lobe involvement between dogs and humans. Indeed, parietal cortex activation was significantly lower in dogs compared to humans during action observation, with most dogs not having any active voxels in the parietal visual and multisensory cortices. Interestingly, although to a lesser degree in humans, we also found contrasting patterns of relative lobe involvement in the two species. Dogs displayed pronounced temporal engagement with significantly more active voxels in the temporal than in the parietal lobe, and task-based connectivity between V1 and the temporal lobe exceeded the connectivity with the parietal lobe. Humans had more active voxels in the parietal than the temporal lobe, but we did not find a difference in task-based functional connectivity between the two lobes. The species’ differences align with their differential pattern of relative lobe expansion^15^. As common marmosets, who also exhibit pronounced temporal activation during action observation^14^, dogs have more expanded occipital and temporal than parietal and frontal lobes. While humans, apes and Old World Monkeys display an opposing trend with significantly more expanded parietal and frontal lobes, rhesus macaques and chimpanzees have stronger frontoparietal activation during action observation^12,13^. Thus, parietal areas may play a more prominent role for action observation in species with more expanded parietal lobes.

The inferior parietal lobule (IPL) – a key parietal region of the human, chimpanzee and macaque AON^12,16,71,72^ – is associated with the planning and execution of reach-to-grasp actions, as well as guiding visual attention (“vision-to-action”; see^73,74^ for reviews), and the anterior supramarginal gyrus portion of the human IPL has been linked to their complex tool-using abilities^75^ (and see^76,77^ for reviews). Another potential explanation for the differential areal activation patterns could, therefore be the species’ tendency for manual object interactions (i.e., with their upper limb). In marmosets, action observation did not elicit IPL activation^14^, and unlike rhesus macaques, chimpanzees or humans, they primarily explore and grasp objects with their mouth rather than their upper limbs^78^. Dogs do not display any tool-using behaviour, and they can only perform reach-to-grasp actions with their snout, which they mainly use to explore their environment. Thus, our findings further elucidate the relationship between the involvement of parietal networks during action observation and the occurrence of complex manual object-manipulating behaviours.

Regarding sensory-motor involvement during action observation, we found partly analogous but also unexpected divergent results. As hypothesized, observing actions performed by conspecifics or heterospecifics elicited activation in human primary and secondary sensory-motor cortices (e.g., inferior frontal gyrus, post-/ precentral gyri^4,17^). In dogs, results also revealed activation in the primary (S1) and secondary (S2) somatosensory cortex (i.e., rostral suprasylvian and ectosylvian gyrus^79,80^). However, despite showing a trend in the direction, S1 activation did not significantly differ from activation for scrambled motion. Contrary to the results in humans, we did not find significant activation in the dog premotor cortex during action observation compared to the control conditions. We only found higher activation for actions compared to both controls in 8 out of the 28 dogs, including mixed-breed and three different pure-breed dogs. It is, therefore, unlikely that specific breeding purposes or behavioural specializations drove the observed differences in activation. It should be noted though that the findings on (pre)motor areas need to be interpreted with caution, as the signal in all dogs’ frontal lobes was affected by their large air-filled sinuses bordering this brain area, resulting in low SNR values and partial up to complete signal drop out in many dogs. Considering this issue and that mirroring activation has been demonstrated in other mammalian and bird species via cell recordings^23–27^, we cannot conclude that dogs do not have premotor areas with mirroring properties. Advances in dog electroencephalography (EEG) research^81–83^, might overcome the limitations of MRI and provide more insights into the involvement of dog sensory-motor and frontal cortices during action observation.

Lastly, due to their close social bond^84,85^, we also explored how dogs and humans perceive each other compared to conspecifics. Interestingly, observing conspecific compared to heterospecific actions elicited increased activation only in human primary visual cortices (V1). However, the reversed contrast led to activation in the entire human AON. Less familiarity with actions performed by hetero-compared to conspecifics and, for humans, uncommon use of the mouth to pick up an object might require increased engagement of the action observation network to understand or predict the observed action, which would align with the predictive coding account of action observation^86^. To the best of our knowledge, only one prior human fMRI study investigated the neural bases while observing actions performed by dogs or humans. However, that study did not directly contrast dog vs. human action^87^. Hence, more neuroimaging research is needed to explore further the neural bases of heterospecific action perception in humans. In dogs, we found no significant activation increases in response to human compared to conspecific actions. The reversed contrast only led to significantly more activation in the mid suprasylvian animate area, which has been previously associated with higher sensitivity towards conspecifics^46^. The lack of pronounced differences between the perception of dog and human actions may be surprising at first sight. Note, however, that all pet dogs participating in our study grew up in human households and are, therefore, highly familiar with human actions, and already dog puppies spontaneously imitate human actions^50^. Thus, dogs’ attention towards human social cues has likely been enhanced through domestication^88^, and their AON is therefore potentially tuned to human and dog actions.

In conclusion, our study marks the first investigation of the dog action observation network. The analogous engagement of somatosensory and temporal areas suggests that both species evolved partly similar networks engaged in action observation. The strong differences in parietal lobe involvement provide exciting new insights into how differential cortical expansions may support brain functions, speaking for the divergent evolution of the species’ object-manipulating behaviours. The findings and our overall approach provide strong foundations and the first stepping stone for future studies investigating the evolutionary history of primate and canine action perception. We hope this will ultimately inform a more advanced understanding of the evolution of the neural bases of social behaviour and learning.

## Acknowledgements

We want to thank Laura Lausegger, Marion Umek and Helena Manzenreiter for training the majority of the dogs and their help collecting the data together with Anna Thallinger, Olaf Borghi and Sara Binder; Boryana Todorova, Marie-Christine Rühle and Theresa Bonkoss for their support in preparing the stimuli set, and all the dogs and their caregivers and human participants for taking part in this project. This project was supported by the Austrian Science Fund (FWF) through project W1262-B29, the Vienna Science and Technology Fund (WWTF) [10.47379/CS18012], the City of Vienna and ithuba Capital AG, the OeAD Marietta Blau grant, and the Messerli Foundation (Sörenberg, Switzerland). The funders had no role in study design, data collection and analysis, decision to publish, or manuscript preparation.

## 4 Author Contributions (CRediT)

**MB**: Conceptualization, Methodology, Software, Formal analysis, Investigation, Data curation, Writing - original draft, Writing – review & editing, Visualization, Project administration. **SK**: Investigation, Writing - review & editing. **ICW**: Conceptualization, Methodology, Writing - review & editing, Supervision. **LLL**: Methodology, Software, Writing - review & editing **LH**: Conceptualization, Resources, Writing - review & editing, Supervision, Funding acquisition. **CL**: Conceptualization, Methodology, Resources, Writing - review & editing, Supervision, Funding acquisition.

## 5 Data availability

The following data is openly available on the Open Science Framework (OSF; osf.io/z479k): Beta maps of all univariate group comparisons, all functionally defined region-of-interest (ROI) masks, detailed sample descriptives, individual motion parameters, raw ROI data (univariate and gPPI), dog whole-brain temporal signal-to-noise ratio maps, and the stimulus material. Raw data is available from the lead author upon request.

## 6 Methods

### 6.1 Participants

#### Action observation task

Twenty-eight pet dogs (*Canis familiaris*; 12 females, age range: 2-12 years, mean age: 6.1 years), extensively trained to undergo MRI scanning^39^, participated in the study. More than half of the sample were pure-bred breeds (9 Border Collies, 5 Australian Shepherds, 1 Labrador Retriever, 1 Nova Scotia Duck Tolling Retriever, 1 Border Collie Australian Shepherd mix, 1 Labrador retriever mix, 1 Small Munsterlander, 1 Leisha dog, 8 mixed breeds). The dogs’ weight ranged from 16-29 kg (mean = 21.58 kg, *SD* = 4.77). The data collection period lasted from May 2020 to March 2023, and we set the minimum sample size to *N* = 12 dogs, which was the median sample size of awake dog fMRI studies at the time of planning this study and still roughly was in 2023 (i.e., by the time of submitting this study). All dogs underwent a veterinary medical check before data collection to assess their general health and eyesight, and the human caregiver gave informed written consent to their participation.

We collected comparative data from *N* = 40 human participants (22 females, age range: 19-28 years, mean age: 23 years) who also participated in an fMRI study investigating face and body perception in dogs and humans^49^ in the same session (data collection period: September to November 2020). We determined the sample size based on previous studies in our lab and other comparative neuroimaging studies (see e.g., Bunford et al., 2020) with similar task designs. Human participants were right-handed with normal or corrected-to-normal vision; they had no history of neurological or psychiatric diseases, did not report fear of dogs and gave informed written consent.

#### Agent localizer

Findings so far about the location of face-and body-sensitive areas in the dog brain were mixed, and a lack of shared template space makes it even more challenging to build on prior work. We therefore additionally used a functional localizer task for the dogs, which allowed us to detect functional analogues of the human ventral temporal pathway (i.e., housing face-and body-sensitive areas^6,8^) within the canine action observation network within our sample. The agent localizer sample comprises *N* = 28 pet dogs (15 females, age range: 2-10, mean age: 5.5 years). Twenty-four of these dogs also participated in the main task on action observation, and data from *n* = 15 dogs was published as part of the original publication of our group investigating agent perception in dogs and human (i.e., Boch et al., 2023).

Detailed sample descriptives are openly available at the study’s OSF data repository. Dog data collection was approved by the institutional ethics and animal welfare commission in accordance with Good Scientific Practice (GSP) guidelines and national legislation at the University of Veterinary Medicine Vienna (ETK-06/06/2017), based on a pilot study conducted at the University of Vienna. The comparative human data collection was approved by the ethics committee of the University of Vienna (reference number: 00565) and performed in line with the latest revision of the Declaration of Helsinki (2013).

### 6.2 Experimental design

#### Action observation task

In two 5-minute task runs, dog and human participants saw videos of a dog or human agent grasping a ball (transitive action) or a video of which the ball was edited out (intransitive action, i.e., identical movement kinematics but no visible action goal). They also saw videos with the agent edited out (object motion condition) to control for object motion with the same trajectory and velocity as in the transitive action video and a phase-scrambled version of the transitive action video serving as a low-level visual and motion characteristics control (see **stimuli section** for details). Thus, participants overall saw six different conditions: dog transitive actions, human transitive actions, dog intransitive actions, human intransitive actions, object motion and phase-scrambled control (see **Figure 1C**). Half of the control condition blocks showed object and scrambled motion based on the human and the other half based on the dog transitive action videos. We presented the videos in a block design (duration: ∼ 12 s; 4 different videos per block), interspersed with a visual baseline (3-7 s jitter) depicting a white cross on grey background using Psychopy^89^. Participants saw three blocks per condition in randomized order in each run, but the same condition was only presented once in a row (i.e., 18 blocks per task run). Video composition for each block and order within blocks were randomized across participants. Dogs and human participants were trained or instructed to attend to the MR compatible screen (32 inch) placed at the end of the scanner bore (passive viewing paradigm). Dogs viewed the task in sphinx position (see **Figure 1A**); to ensure they could see the videos without looking up or moving their head, we presented the videos and implicit baseline at their eye level (i.e., 100 pixels below the centre of the screen).

#### Agent perception localizer

In two 5-minute task runs, dogs saw colour images of faces and (headless) bodies of dogs and humans, as well as inanimate objects and scrambled images (i.e., low-level visual control) presented on a grey background. As in the main task, images were presented in a block design (12 s blocks, 5 images/block) with the image composition randomized and no consecutive blocks of the same stimulus conditions (see^49^ for details on the paradigm and stimulus material).

#### Procedure

Before data collection, dogs received extensive training to habituate to the scanner environment to participate without any sedation or restraints (see^39^ for a detailed description of the training and data collection procedure). A trainer was also present in the scanner room to monitor and handle the dog out of its sight. We positioned the camera of an eye-tracker (Eyelink 1000 Plus, SR Research, Ontario, Canada) below the MRI screen to live-monitor participants’ attention towards the screen (i.e., whether eyes were open and directed towards the screen) and motion.

Dogs and humans were equipped with earplugs, and both could stop data collection anytime, either by leaving the scanner by retreating from the coil and exiting the scanner via a custom-made ramp (dogs) or pressing an alarm squeeze ball signalling to stop the scanning (humans). Human participants completed both task runs within one session with a short break in between. After each session, we evaluated the motion parameters. If overall motion for any of the three translation directions exceeded 4 mm or if scan-to-scan motion (i.e., framewise displacement to account for translational and rotational motion^90,91^) exceeded .5 mm in more than 50% of the scans, we re-invited the dog to repeat the task run (individual motion parameters are available on the study’s OSF data repository). Based on these criteria, and sufficient attentiveness evaluated by the research team based on live video observation, dog participants needed on average 2.75 sessions (*SD* = 1.46) to complete both task runs successfully. Dogs had at least a one-week break in-between data collection sessions.

#### Stimuli

All stimuli were created based on video recordings of a dog or human agent picking up different toy balls (see **Figure 1B-C** for examples). To increase ecological validity and familiarity with the observed action, we asked human agents to grasp the ball with their (right) hand. We trained the dog agents to pick it up with their mouth as this would be their natural way to perform this task. We recruited two human and dog models (each a male and a female) and recorded four videos from each agent (duration: ∼3 s). Videos were filmed in the same setting to ensure no differences in lighting between videos, and human and dog models were unfamiliar to the study participants. The original videos were cropped to 720 × 720 pixels, and shadows and the dog trainer were edited out to create the transitive action video. We created the intransitive action stimuli by editing out the ball from the transitive action video, the object motion control by editing out the agent and finally, the scrambled low-level visual control by phase-scrambling the transient action video (see **Figure 1C**).

### 6.3 MRI data acquisition

MRI data for the action observation task was obtained using a custom-made 16- channel (k9) head coil for the dog participants^53^ and a 32-channel head coil for human participants, used in a 3T Siemens Skyra MR-system (Siemens Medical, Erlangen, Germany). For three dogs, we used structural scans which had been previously acquired with a 15-channel human knee coil due to unavailability of the dogs to acquire a new structural scan. However, structural sequences for both coils were the same and qualitative comparisons showed that image registration worked equally well with structural scans acquired from both coils. For the agent localizer, data from *n* = 15 dogs (i.e., from the original publication of the localizer paradigm^92^) were acquired with the human knee coil and from *n* = 13 dogs with the k9 head coil.

For all dog functional scans, we used a 2-fold multiband (MB) accelerated echo planar imaging (EPI) sequence for the dog functional scans with the following parameters: voxel size = 1.5 × 1.5 × 2 mm^3^, repetition time (TR) / echo time (TE) = 1000/38 ms, field of view (FoV) = 144 × 144 × 58 mm^3^, flip angle = 61°, 20% gap and 24 axial slices (interleaved acquisition, descending order). Individual numbers of volumes per action observation task run vary slightly because task acquisition was stopped manually (mean = 324 volumes; *SD* = 10). Voxel size for all structural scans was .7 mm isotropic (TR/TE = 2100/3.13 ms, FoV = 230 × 230 × 165 mm^3^).

We acquired human functional scans (each run: mean = 272 volumes *SD* = 2) with a 4-fold MB accelerated EPI sequence: voxel size = 2 mm isotropic, TR/TE = 1200/34 ms, FoV = 192 × 192 × 124.8 mm^3^, flip angle = 66°, 20% gap and 52 axial slices coplanar to the connecting line between anterior and posterior commissure (interleaved acquisition, ascending order). In the same orientation as functional scans, we obtained additional field map scans to correct for magnetic field inhomogeneities using a double echo gradient echo sequence with a voxel size of 1.72 x 1.72 x 3.85 mm^3^, TR/TE1/TE2 = 400/4.92/7.38 ms, FoV = 220 × 220 × 138 mm^3^, flip angle = 60° containing 36 axial slices. Voxel size for structural scans was .8 mm isotropic with TR/TE = 2300/2.43 ms and FoV = 256 × 256 × 166 mm^3^.

### 6.4 Data preprocessing

We preprocessed and analyzed imaging data of both species using SPM12 (https://www.fil.ion.ucl.ac.uk/spm/software/spm12/), Matlab 2020b (MathWorks Inc., MA, USA) and R 4.3.0^93^. For the dogs, after slice timing correction (reference: middle slice) and realignment, we manually reoriented the functional images and set the origin at the anterior commissure using the SPM *Display* function to match the orientation of the structural images and the dog template^55^. Next, we skull-stripped the structural images using individual binary brain masks created with itk-SNAP^94^, co-registered the brain-extracted structural images to the mean functional images, and segmented them. We then normalized and resliced (1.5 mm isotropic) all imaging data to the breed-averaged dog template space^55^ and smoothed the data with a 3-dimensional Gaussian Kernel (full-width-at-half-maximum, FWHM; 3 mm; i.e., twice the raw within-plane voxel resolution).

For the human data, we realigned and unwarped functional images after slice time correction (reference: middle slice) using the individual field maps. Individual structural images were then co-registered to the mean functional images and segmented. Next, we normalized and resliced (1.5 mm isotropic) the imaging data and applied spatial smoothing with a 3-dimensional Gaussian Kernel (FWHM, 4 mm; i.e., twice the raw voxel resolution).

Next, we calculated individual scan-to-scan motion (i.e., framewise displacement, FD) to account for translational and rotational motion in both species. For each scan exceeding the a priori set FD threshold of .5 mm, we added a motion regressor to the first-level general linear models (GLMs; motion scrubbing, Power et al., 2012, 2014). On average, 5.7% of the dog scans (run 1: mean FD = .2 mm, 90^th^ percentile = .35 mm; run 2: mean FD = .21 mm, 90^th^ percentile = .32 mm) and 1.5% of the human scans (run 1: mean FD = .18 mm, 90^th^ percentile = .23 mm; run 2: mean FD = .18 mm, 90^th^ percentile = .25 mm) exceeded the threshold in each run.

### 6.5 Overview of the main analyses and rationale

The aim of the first part of our analyses was to localize the dog action observation network for the first time and, based on that, to identify functional analogies between the dog and human action observation networks by applying whole-brain univariate and region-of-interest (ROI) analyses. To this end, we applied the following analysis steps.

First, we identified the species’ action observation networks by comparing activation for both agents and action types (i.e., pooled activation) to the implicit baseline and the two control conditions. Second, we tested our hypothesis of comparable activation for transitive vs. intransitive actions due to the species’ imitation skill, and we explored how the human and dog action observation network responded to observing actions performed by conspecifics vs. heterospecifics. Third, due to anticipated low tSNR in parts of the sensorimotor cortices, we also conducted a univariate ROI analysis in addition to the whole-brain analysis to test our hypothesis of somatosensory and (pre-) motor activation in response to action observation as this approach results in higher sensitivity. Fourth, we used a functional localizer task to detect face-and body-sensitive areas (i.e., agent areas) in the dog temporal lobe, which allowed us to identify the functional analogues of the human ventral visual pathway^8,95^ within the dog action observation network and to test our hypothesis of temporal lobe engagement beyond the agent areas in dogs, functionally analogous to humans (i.e., lateral temporal or third visual pathway^6,7^).

The second part of our analysis then focused on between-species divergencies. Here, we tested our key hypothesis of functional divergencies in temporal vs. parietal lobe engagement in the two species. Based on prior non-human primate and human research and the species’ differential relative lobe expansion, we specifically expected a stronger temporal than parietal lobe engagement during action observation in dogs but a more balanced involvement of both lobes in humans. We tested this hypothesis by implementing two analysis approaches. First, we conducted within-and cross-species comparisons of the extent of active voxels in each species’ temporal and parietal lobe during action observation. Second, we compared task-based functional connectivity with the primary visual cortex and the temporal vs. parietal lobe. This also allowed us to explore information exchange within the species’ action observation networks.

### 6.6 Univariate analysis

#### 6.6.1 Action observation task

##### First-level analysis

First-level analyses were carried out using a GLM approach implemented in SPM12 (https://www.fil.ion.ucl.ac.uk/spm/software/spm12/). We defined four task regressors for our main conditions of interest: transitive and intransitive actions of each species (i.e., dog/human × transitive/intransitive action) and two for the control conditions (i.e., object motion and scrambled). All blocks were estimated as a boxcar function time-locked to the onset of each block with a duration of 12 s and convolved with a tailored dog haemodynamic response function^52^ (HRF) or the standard human HRF implemented in SPM12. In addition, we added the six motion regressors retrieved from image realignment and the individual framewise displacement regressors as nuisance regressors. We applied a temporal high-pass filter with a cut-off at 128 s. Due to signal distortions caused by dogs’ large sinuses potentially affecting signal in the dog frontal lobes and, therefore, partly the sensory-motor cortices (see **Supplementary Figure S1B-C**), we applied implicit masking to exclude voxels with a mean value lower than 80% of the global signal. We also calculated individual whole-brain temporal signal-to-noise ratio (tSNR) maps 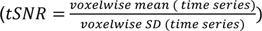 to measure where large sinuses might have affected individual brain signal in dogs.

Finally, we computed contrasts for all six task regressors, an action observation contrast averaging the activation for transitive and intransitive actions of both species (i.e., all conditions showing agent performing an action > implicit visual baseline) and a task-activation contrasts (i.e., all task conditions > implicit visual baseline).

##### Whole-brain group comparisons

First, we conducted a one-sample *t*-test for the action observation contrast (i.e., all action conditions > implicit visual baseline) for a first exploration of the action observation network and one sample *t*-tests for each condition of interest. We then implemented two paired-sample *t*-tests to compare the activation elicited by action observation to (1) the low-level visual stimulation control (i.e., all action conditions > phase-scrambled motion) and (2) the activation elicited by object motion (i.e., all action conditions > object motion condition). To investigate if the dog and human action networks respond to conspecific and heterospecific actions and to test our hypothesis of no pronounced differences in activation for transitive and intransitive actions due to the species imitation skills, we calculated a within-subjects full factorial model using the flexible factorial framework in SPM12 with the factors *action* (transitive, intransitive) and *agent* (dog, human) to test for main effects of action and agent, and an action × agent interaction.

For all whole-brain group analyses, we determined significance by applying cluster-level inference with a cluster-defining threshold of *p* < .005/.001 (dogs/humans) and a cluster probability of *p* < .05 family-wise error (FWE) corrected for multiple comparisons. We chose a lower cluster-defining threshold for the dog data as this has been commonly used in the field of dog fMRI (see e.g.^46,49,63^), suggesting its effectiveness in identifying relevant brain responses while at the same time increasing comparability to and integration of the present data with prior findings data. We derived the cluster extent (i.e., the minimum spatial extent to be labelled significant) using the SPM extension “CorrClusTh.m”^96^. Anatomical labelling of activation peaks and clusters of all reported results refers to the dog brain atlas from Czeibert and colleagues^54^ normalized to the stereotaxic breed-averaged template space^55^, and the Harvard-Oxford human brain atlas^56^; and was performed using the python software AtlasReader^97^.

##### Univariate region-of-interest (ROI) analysis of dog sensorimotor cortex

We hypothesized somatosensory and (pre-) motor areas to be part of the action observation network in dogs, functionally analogous to humans. Due to the anticipated lower tSNR in this area in dogs, we conducted an ROI analysis to test this hypothesis with higher sensitivity. Constrained masks for somatosensory and motor cortices of the dog brain do not exist, and knowledge about the exact locations, especially of pre-and secondary motor cortices, is limited. However, it is known which gyri house sensorimotor cortices. The premotor cortex and supplementary motor area (SMA) are located in the precruciate gyrus. The motor cortex is housed in the anterior portion of the postcruciate gyrus, bordering the primary somatosensory cortex (S1) posteriorly at the postcruciate sulcus and ventrally at the rostral suprasylvian sulcus^98,99^. S1 encompasses the posterior postcruciate gyrus and the rostral suprasylvian gyrus, and the secondary somatosensory cortex (S2) is housed in the rostral ectosylvian sulcus^79,80^; see also **Supplementary Figure 1A**). We, therefore, defined four ROIs using the following publicly available gyri masks^54^: precruciate gyrus (“premotor and SMA”), postcruciate gyrus (“M1/S1” or “somatomotor”), rostral suprasylvian gyrus (“rostral suprasylvian S1”), rostral ectosylvian gyrus (“S2”). To constrain the large anatomical masks, we created a binary mask of the group task-based activation (i.e., all conditions > implicit visual baseline) liberally thresholded at *p* = .05 uncorrected and intersected the functional mask and each of the anatomical gyri masks. Next, using the REX toolbox^100^, we extracted parameter estimates for each condition-of-interest (i.e., pooled actions contrast, dog/human transitive/intransitive action) and the controls (i.e., object and scrambled motion) from the sensorimotor ROI masks.

First, to test whether areas of the sensorimotor cortex respond stronger to action observation compared to controls, we employed linear mixed models (LMMs) for each of the four ROIs using the R packages *lme4*^101^ and *afex*^102^. We defined activation levels (i.e., parameter estimates) as the dependent variable, *condition* (levels: actions, object motion, phase-scrambled motions) as predictor, and added per-subject random intercepts. To investigate potential differences in activation due to the observed agent or action type, we conducted LMMs for each ROI, with activation levels as the dependent variable and *agent* (levels: dog, human) and *action* (levels: transitive, intransitive) as predictors (i.e., 2 × 2 within-subjects design) and added by-subject random intercepts. We applied false-discovery rate (FDR) control to correct *p*-values for group comparisons investigating the same research questions (e.g., greater activation for action observation compared to controls) and for planned post hoc comparison.

##### Cross-species comparison of parietal and temporal cortex involvement

As outlined above, we predicted stronger temporal than parietal lobe activation in dogs and significantly less activation in the dog compared to the human parietal lobe during action observation. To quantitatively test the potential cross-species differences in parietal and temporal activation, we measured the individual percentages of active voxels within each lobe during action observation (i.e., 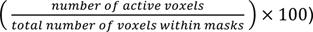. Specifically, we focused on the visual and multisensory cortices and excluded somatosensory and auditory areas of both species from the lobe masks and subsequent analysis.

For the dogs, we created the parietal lobe mask by combining bilateral anatomical masks from Czeibert and colleagues^54^ of the anterior portions of the marginal and ectomarginal gyrus with the ascending ramus indicating the posterior border, and the presplenial gyrus^103^. Dog temporal lobe definitions vary regarding the in-or exclusion of the mid and caudal suprasylvian gyrus^55,103^. Considering the functional convergence between the mid and caudal suprasylvian agent areas and the human inferior temporal cortex^46,49^, we included both gyri in the dog temporal lobe mask together with the caudal composite and rostral sylvian gyri (see **Supplementary Figure S2** for main gyri of the dog brain and masks). We created the human lobe masks using anatomical masks from the Harvard-Oxford brain atlas^56^. The bilateral temporal mask includes the temporal pole, all middle and inferior temporal gyrus masks and the temporal fusiform cortex. For the parietal lobe mask, we combined bilateral masks of the superior parietal lobule, supramarginal gyrus (anterior and posterior division), angular gyrus and precuneus cortex.

As for the group analysis, we determined significantly active voxels, by thresholding the individual contrast maps for the action observation contrast (i.e., all conditions displaying an agent performing an action > implicit baseline) by applying cluster-level inference. We also conducted secondary analyses with more liberal thresholds to ensure cross-species comparisons were not biased due to too conservative thresholds. First, we applied an uncorrected threshold of *p* < .005/.001 for dogs/humans) to determine significant voxels. Second, following procedures established in comparative primate neuroimaging research ^12,104^, we defined the most active voxels (i.e., highest positive beta-values) as significant. We used the top 5% voxels as the threshold but also calculated parietal and temporal cortex involvement for percentage thresholds ranging from 1% to 100% in steps of 5% for a further visual inspection.

For group (cross-species) comparisons, we employed a LMM with proportions (i.e., individual percentages of active voxels) as the dependent variable and *lobe* (levels: parietal, temporal; within-subject) and *sample* (levels: dog, human participants; between-subject) as predictors and added by-subject random intercepts. *P*-values for planned post hoc comparisons were FDR-controlled.

#### 6.6.2 Agent localizer

We employed a functional localizer task to locate face-and body-sensitive regions (i.e., agent regions) in our sample in order to identify the functional analogues of the human ventral visual pathway^8^ within the dog’s action observation network.

##### First-level analysis

As described above, first-level analyses were conducted using a GLM approach implemented in SPM12. We defined four task regressors for the main conditions of interest: faces and bodies of each species (i.e., dog/human faces/bodies) and two for the control conditions (i.e., inanimate objects and scrambled). We then computed averaged contrasts for faces and bodies (i.e., pooled activation for dog and human stimuli), for dogs and humans (i.e., pooled for faces and bodies), and for inanimate objects (all conditions > scrambled control).

##### Group comparison

To localize the face and body-sensitive areas, we conducted a whole-brain one-way repeated measures analysis of variance (ANOVA; levels: faces, bodies, inanimate objects; all conditions > scrambled controls) and a second ANOVA to investigate potential differences due to species depicted with the factors *agent* (dog, human) and *section* (face, body; all conditions > scrambled controls). Both ANOVAs were implemented using the flexible factorial framework in SPM12.

### 6.7 Task-based functional connectivity analysis

As outlined above, we expected stronger temporal than parietal lobe engagement during action observation in dogs, but a more balanced involvement of the two lobes in humans. In addition to the comparison of activation extent, we also aimed to compare the strength of information exchange (i.e., task-based functional connectivity) with V1 between the parietal and temporal lobe in the two species.

#### 6.7.1 Action observation task

##### First-level analyses

To investigate if action observation led to differences in functional connectivity of the primary visual cortex (i.e., seed region) with the temporal and parietal cortices in both species, we used generalized psychophysiological interaction (gPPI) analyses^40^. For the dogs, due to the unavailability of an anatomical mask, we created a primary visual cortex (V1) sphere (x = 1, y = -29, z = 16, 4 mm) using coordinates from a visual localizer task^52^. We built the human V1 seed region by combining the supra-and intracalcarine cortex masks from the Harvard-Oxford brain atlas^56^. From the respective V1 masks, we extracted the first eigenvariate of the individual functional time courses of the dog and human participants, adjusted the functional time courses for average activation using an F-contrast, and deconvolved them to estimate the neural activity in the seed region (i.e., physiological factor). We then multiplied the neural activation estimate with a boxcar function time-linked to the onset of each block (i.e., psychological factor) convolved with the species-specific HRF models^52^. This resulted in one psychophysiological interaction regressor per condition of interest. The interaction regressors were added to the first-level design matrix, and the GLMs were estimated. Using the REX toolbox^100^, we then extracted mean functional connectivity estimates between the seed region (i.e., V1) and agent-and action-sensitive areas in both species (i.e., target regions) for each condition-of-interest (i.e., pooled actions contrast, dog/human transitive/intransitive action) and the controls (i.e., object and scrambled motion).

##### Group comparisons

On the group level, we investigated whether action observation led to greater functional connectivity between V1 and agent-and action-sensitive areas than the control conditions and whether task-based functional connectivity between V1 and the temporal vs. parietal lobe differed in dogs and humans (see also section **Regions-of-interest task-based functional connectivity** below). First, we aggregated the functional connectivity estimates across all anatomical masks of the same lobe and employed an LMM for each species with the aggregated functional connectivity as the dependent variable, *lobe* (levels: parietal, temporal; within-subject) and *condition* (levels: actions, object motion, scrambled motion) as predictors and random intercepts as well as random slopes for *lobe* and *condition*. This corresponds to the maximal random effects structure possible with this design, as recommended by^105^. The factor *condition* was contrast coded using Helmert coding (contrast 1: actions – (object motion + scrambled motion) / 2; contrast 2: scrambled motion – object motion) to directly test our contrast of interest (i.e., contrast 1). Finally, we investigated task-based functional connectivity changes in each anatomical area separately by setting an LMM with functional connectivity as the dependent variable for each anatomical area, *condition* (levels: actions, object motion, scrambled motion) as the predictor, and we added by-subject random intercepts. *P*-values for planned post hoc comparisons and analyses investigating the same research question were FDR-controlled.

##### Regions-of-interest task-based functional connectivity

The anatomical areas included in the parietal and temporal lobe masks described above (section **Cross-species comparison of parietal and temporal cortex involvement**) served as the target regions for the dogs. For the human participants, we used anatomical masks (retrieved again from the Harvard-Oxford brain atlas^56^) of the known temporal and parietal core nodes of the action observation network. This included the superior parietal lobule, the supramarginal gyrus in the human parietal lobe, and the posterior fusiform cortex, and the posterior superior temporal sulcus (pSTS) area in the temporal lobe. Due to the lack of a pSTS mask in volumetric space, we created and subsequently combined left and right hemisphere spheres based on coordinates from previous work investigating the functional organization of the STS^106^ (sphere radius: 10 mm; left centre x = -50, y = -48, z = 15; right: x = 50, y = -47, z = 13). Finally, we removed all voxels overlapping with the pSTS mask from the supramarginal gyrus mask.

Our rationale to define the target regions for the main functional connectivity analysis anatomically was because the univariate activation analysis did not reveal any action-sensitive areas in the dog parietal lobe, and we did not want the lobe functional connectivity comparison to be biased by comparing more constrained functionally defined target areas with anatomical masks. However, in a secondary analysis, we also investigated functional connectivity between V1 and functionally defined action-and agent-sensitive areas in the dog temporal lobe based on the univariate activation results (i.e., face-and body-sensitive areas localized with the agent-localizer and cluster resulting from the overlap actions > object motion ∩ actions > scrambled motion) to asses connectivity of the areas identified as part of the dog action observation network.

#### 6.7.2 Agent localizer

The ventral visual pathway expands from the primary visual cortex to the inferior temporal cortex of humans ^8,95^, representing one component of the human action observation network. The univariate analysis of the functional (agent) localizer served to detect the functional analogues of this pathway within the dog action observation network. In this secondary analysis, we further examined if the localized dog face-and body-sensitive areas are functional analogues of the human ventral visual pathway^8,95^ by investigating if these areas exchange information with V1 during the perception of static faces and bodies compared to controls.

##### First-level analysis

We employed gPPI analyses as described above in detail using the same V1 sphere. Using the REX toolbox^100^, we then extracted mean functional connectivity between the seed region and agent-sensitive areas in dogs (i.e., target regions) for each condition-of-interest (i.e., faces, bodies, dog/human faces/bodies) and the controls (i.e., inanimate objects, scrambled images). Target regions in the mid and caudal suprasylvian gyrus were defined based on the univariate results from the contrasts faces > controls, bodies > controls, bodies > faces.

##### Group comparison

To investigate task-based functional connectivity changes between V1 and the agent areas, we set up one LMMs for each region with functional connectivity as the dependent variable. We defined *condition* (levels: faces, bodies, inanimate objects, scrambled images) as the predictor and added by-subject random intercepts. *P*-values for planned post hoc comparisons and analyses investigating the same research question were FDR-controlled.

