## Supplementary material for "Action observation reveals a network with divergent temporal and parietal lobe engagement in dogs compared to humans"

#These authors share senior authorship.

### 1 Supplementary figures

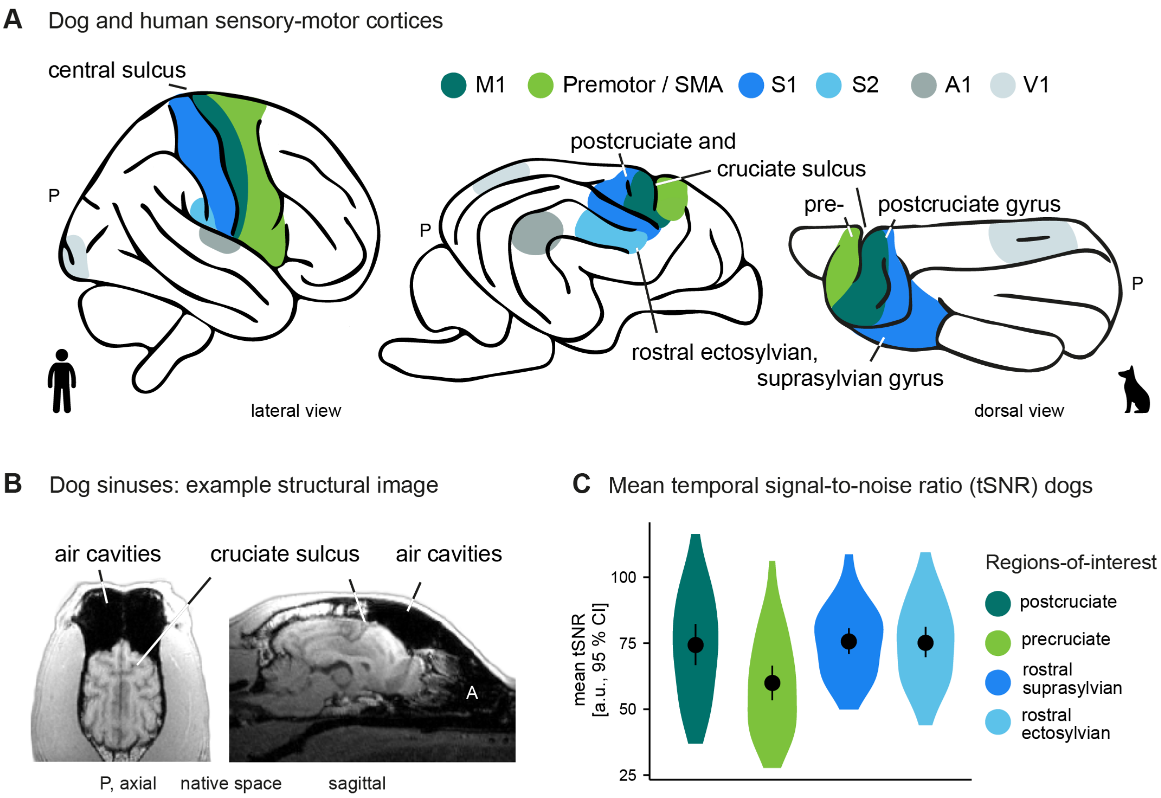


**Figure S1**. **Decreased temporal signal-to-noise (tSNR) ratio in dog pre- and secondary motor cortices housed in precruciate gyrus**. (**A**) For visual comparison and guidance, we created a schematic figure of the human and dog sensory-motor cortices along with the primary visual (V1) and auditory (A1) cortices; see also **Supplementary figure S2** for a schematic overview of all dog gyri and sulci mentioned in the study. The central sulcus marks the border between primary somatosensory (S1) and motor (M1) cortices in humans. The dog analogue of the human central sulcus is the less pronounced postcruciate sulcus, indicating the transition between M1 and S cortices. S1 further extends to the dog rostral suprasylvian gyrus, and S2 is located in the rostral ectosylvian gyrus ventral to S1. The cruciate sulcus marks the transition from M1to pre- and supplementary motor (SMA) cortices housed in the precruciate gyrus[1–4]. (**B**) As presented in the example structural image from one of the dogs in our study sample (Border Collie Australian Shepherd mix), dogs have large sinuses locate anterior-superior to the frontal lobes, heavily affecting the signal and resulting in signal drop out in these areas when using T2*-weighted (fMRI) scans. This includes the precruciate gyrus in large parts (i.e., dog pre- and supplementary motor areas). (**C**) Temporal signal-to-noise ratio (tSNR) measures were considerably lower in the precruciate compared to the other gyri housing sensor-motor areas. The violin plots show the group mean tSNR measured in arbitrary units (a.u.) with error bars indicating the 95% confidence interval (CI) and kernel density plots. Individual whole-brain tSNR maps are provided on the study's OSF data repository. P, posterior. The dog and human icons in A were purchased from thenounproject.com (royalty-free license).

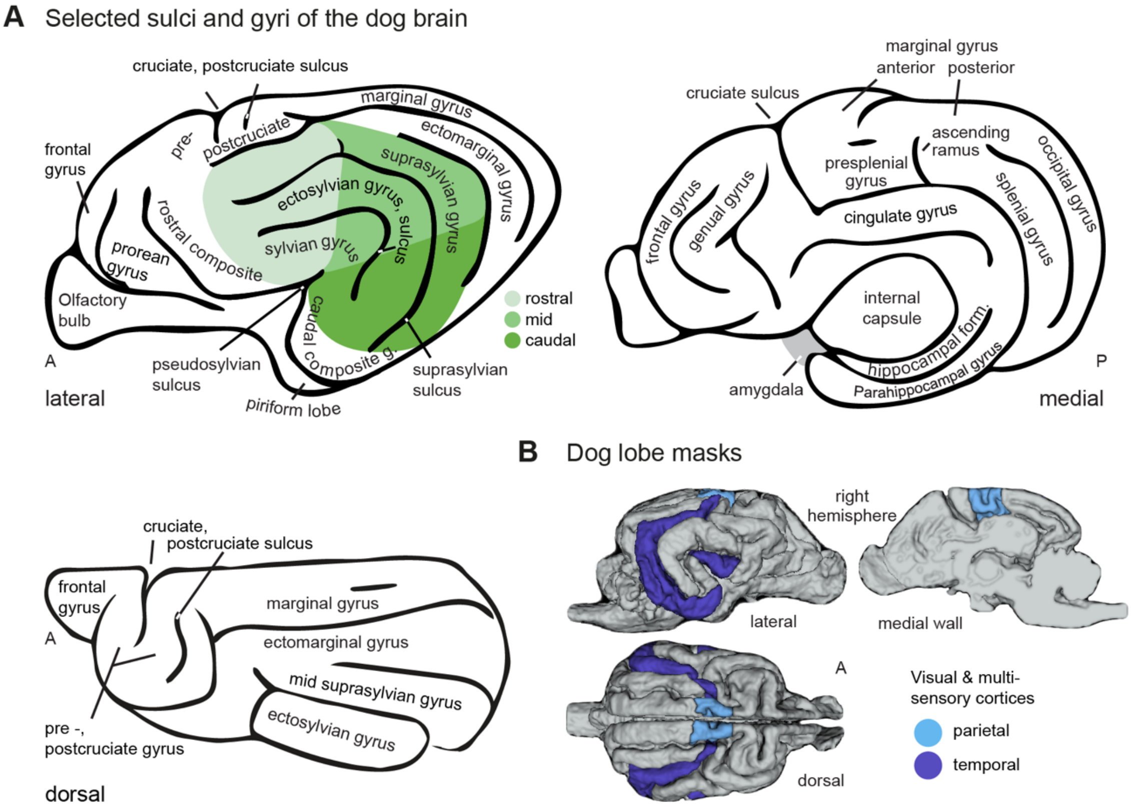


**Figure S2. Selected gyri and sulci of the dog brain and masks used for the temporal and parietal lobes.** (**A**) Lateral, medial and dorsal view of the dog brain (schematic drawings), including relevant gyri and sulci for the present study, accompanied by major anatomical landmarks for visual guidance. (**B**) The lobe masks included visual and multisensory areas of dogs' parietal (anterior marginal and ectomarginal, and presplenial gyrus) and temporal (mid and caudal suprasylvian, rostral sylvian and caudal composite gyrus) cortices. Auditory (temporal caudal and mid ectosylvian gyrus and caudal sylvian gyrus) and somatosensory cortices (parietal rostral suprasylvian gyrus and postcruciate gyrus) were excluded. A, anterior; P, posterior

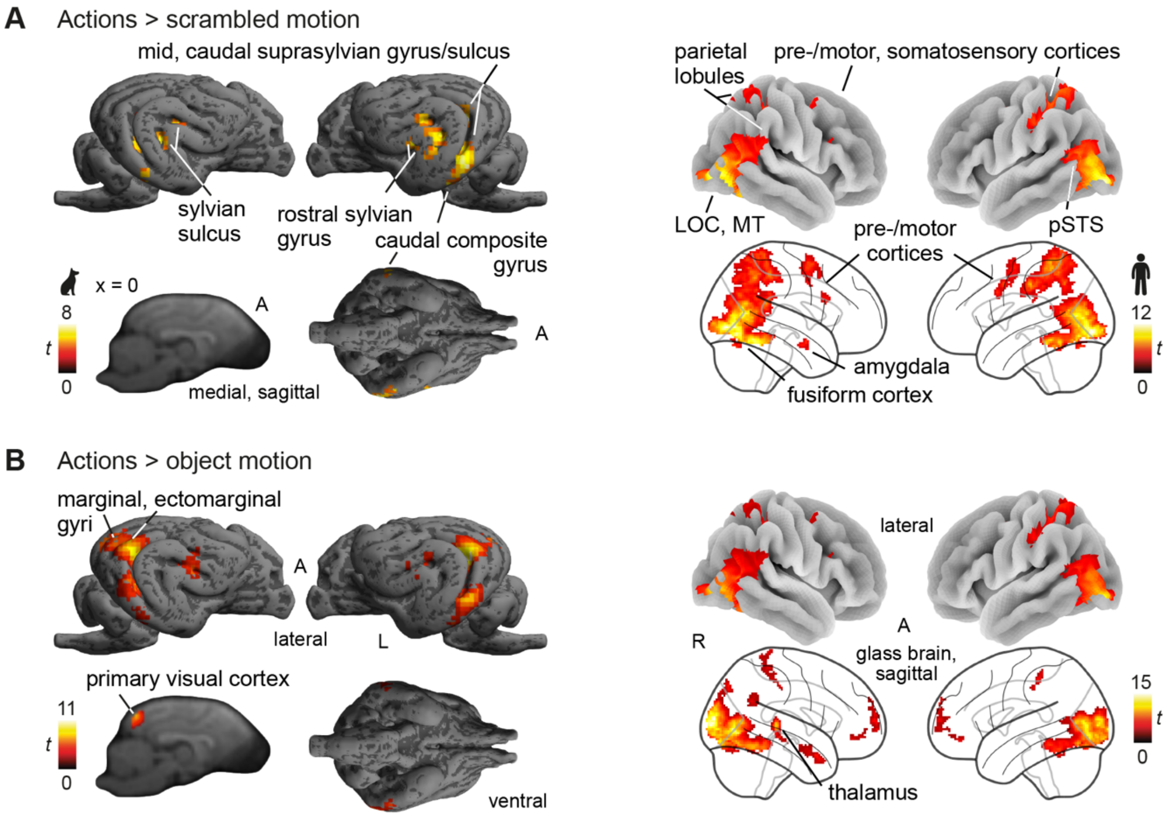


**Figure S3**. **Action observation compared to object and scrambled controls in the dog and human brain.** (**A**) Observing actions (i.e., pooled activation for transitive and intransitive actions performed by dogs and humans) contrasted with scrambled low-level visual control stimuli elicited activation in the dog and human occipito-temporal lobe. In humans, we additionally observed parietal, premotor and somatosensory activation. (**B**) Action observation compared to object motion also led to greater activation in occipito-temporal cortices of both species, but also in the primary visual cortex. Parietal activation was exclusive to humans; we did not find premotor activation in either species. Results are $p < .05$ FWE-corrected at cluster-level using a cluster-defining threshold of $p < .005/.001$ (dogs/humans); see also **Supplementary Tables S1-S2**. A, anterior; L, left; R, right; *t*, *t*-values. The dog and human icons in A were purchased from thenounproject.com (royalty-free license).

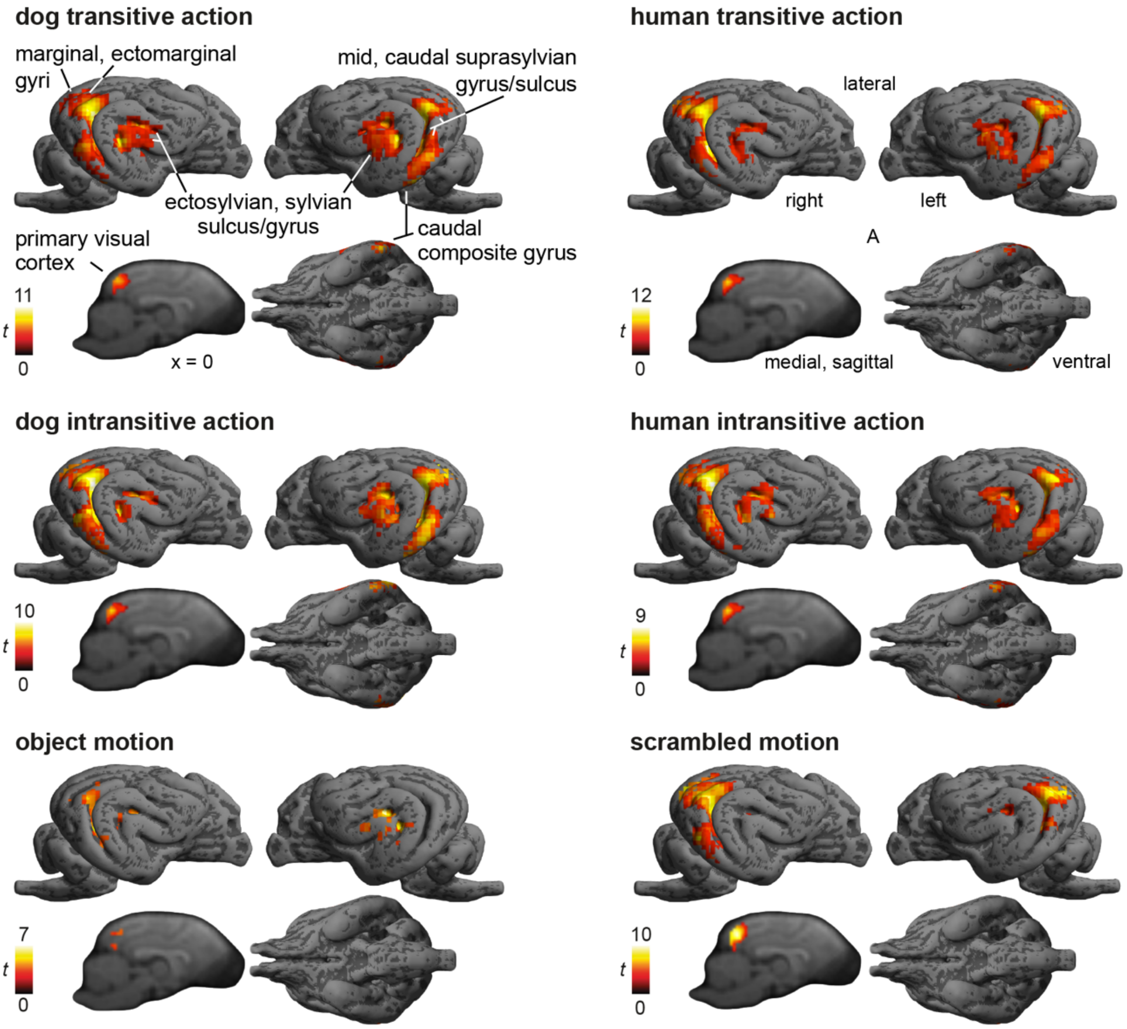


**Figure S4. Functional maps of each condition-of-interest compared to implicit visual baseline (dogs)**. For comparability, activation for each condition is displayed on left and right lateral and ventral surface renders, and a sagittal section (x = 0) to show primary visual cortex activation. Anatomical labels and locations described in the first row therefore apply to all contrast images. Results are $p < .05$ FWE-corrected at cluster-level using a cluster-defining threshold of $p < .005$. A, anterior; *t*, *t*-values.

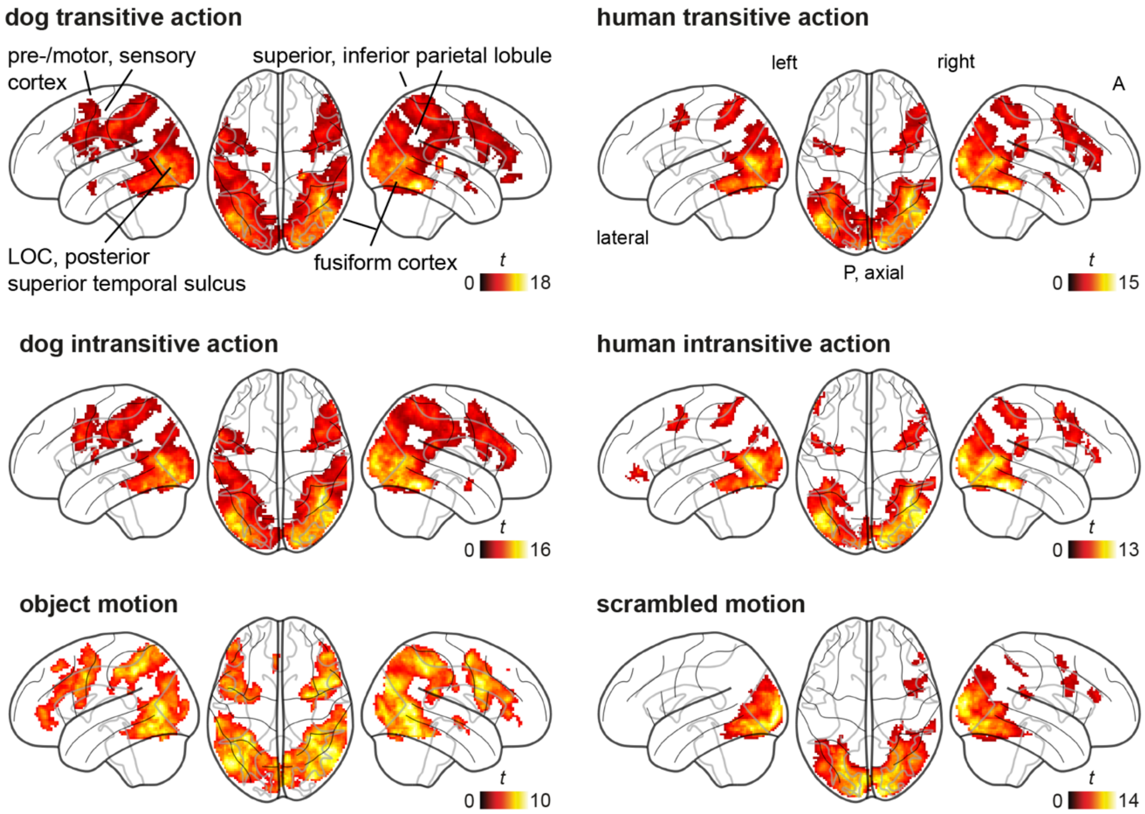


**Figure S5. Functional maps of each condition-of-interest compared to implicit visual baseline (humans)**. Anatomical labels and locations described in the first row apply to all contrast images. Results are $p < .05$ FWE-corrected at cluster-level using a cluster-defining threshold of $p < .001$. A, anterior; P, posterior; LOC, lateral occipital cortex; *t*, *t*-values

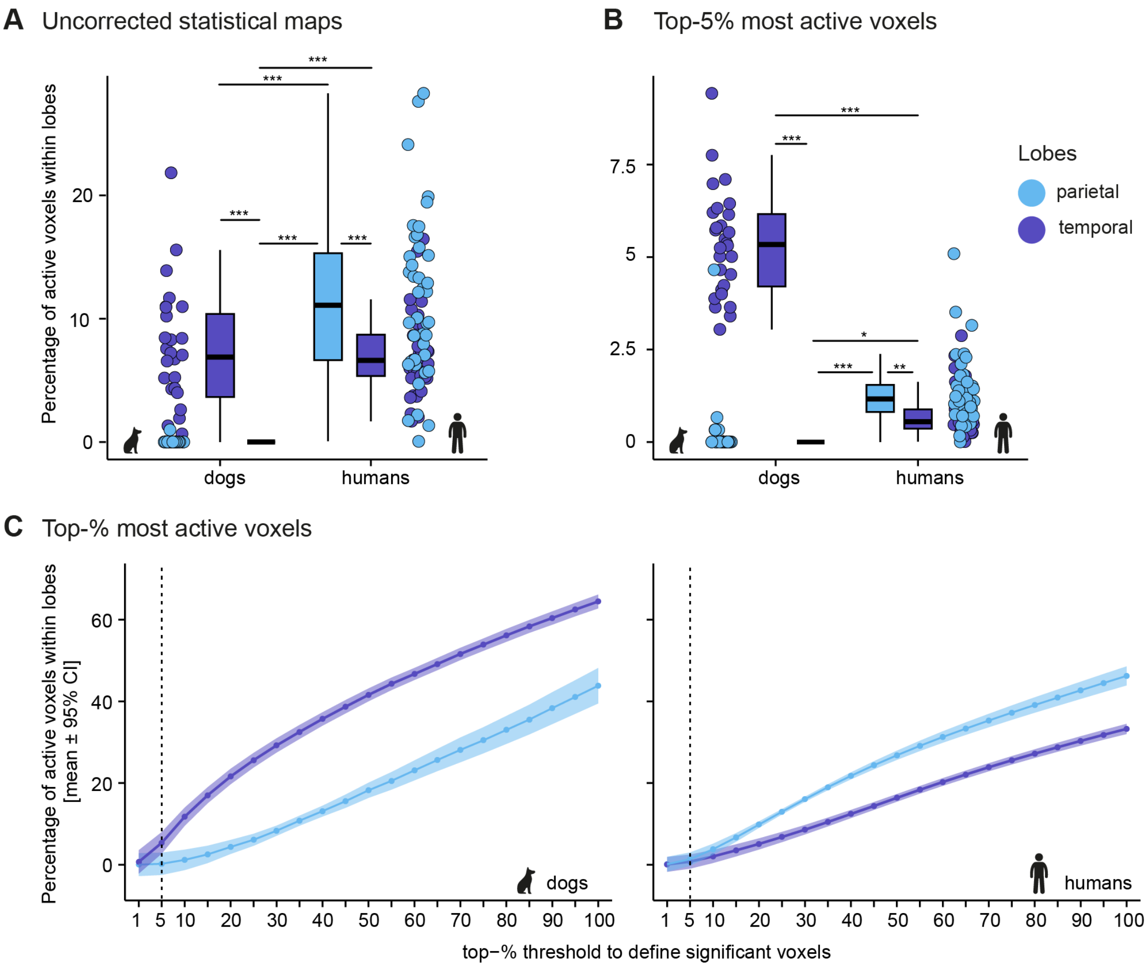


**Figure S6. Secondary analysis with more liberal thresholds to determine active voxels confirms stronger temporal engagement in dogs.** To ensure that cross-species comparisons were not affected by differences in power, we also applied more liberal thresholds to determine individual active voxels during action observation compared to implicit visual baseline. (**A**) First, we classified all voxels surviving the uncorrected cluster-threshold of $p < .005/001$ (dogs/humans) as active voxels. (**B**) Second, we chose an even more liberal threshold and defined the 5% voxels with the highest activation levels as significant. (**A**,**B**) Results for both analyses confirmed the findings of the main analysis with higher proportions of active voxels in the human compared to the dog parietal lobe and a significant temporal activation bias in the dog brain, and significantly more active voxels in the human parietal compared to the temporal lobe (see also **Supplementary Table S6**). The boxplots show the median percentage (black horizontal line), interquartile range (box), the lower/upper adjacent values (whiskers), and are accompanied by coloured dots representing the individual percentages. (**C**) Finally, since top-5% was an a priori but arbitrary threshold to determine active voxels, we also explored proportional lobe engagement for thresholds ranging from the top-1% most active to including all voxels with higher values compared to baseline (i.e., 100%) in steps of 5% (x-axis). These plots illustrate that both species showed reversed parietal vs. temporal involvement (dogs: temporal > parietal, humans: parietal > temporal) and that this is not affected by the %-threshold; dogs, however, show a more pronounced discrepancy between lobes. The plots indicate the mean percentage of active voxels for each threshold (coloured dots) with error bars indicating the 95% confidence interval (CI; ribbon). Planned comparisons were false discovery rate (FDR) corrected to control for multiple comparisons * $p < .05$, ** $p < .01$, ** $p < .001$. The dog and human icons were purchased from thenounproject.com (royalty-free license).

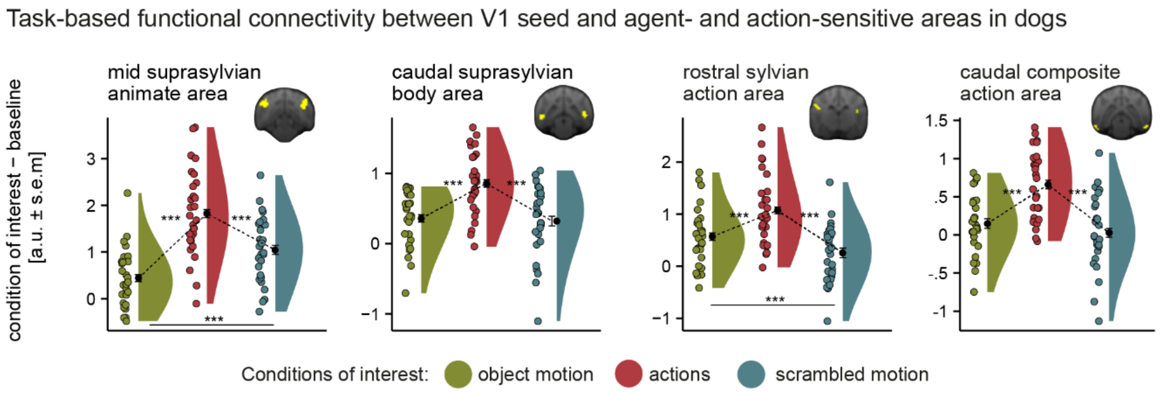


**Figure S7. Strongest task-based functional connectivity between primary visual cortex and temporal agent and action areas during action observation.** Action observation led to the highest connectivity between each area and the primary visual cortex (V1) during action observation when compared to both control conditions. V1 connectivity was significantly higher for object than scrambled motion in the caudal suprasylvian body and composite action areas (see **Supplementary Table S10**). Regions-of-interest (ROIs) were functionally defined based on the univariate results from the action observation task (i.e., caudal composite and rostral sylvian action areas, **Figure 3B**) and the agent localizer (i.e., mid suprasylvian animate and caudal suprasylvian body area, **Figure 3C**). Voxels overlapping with the rostral ectosylvian gyrus (i.e., secondary somatosensory cortex) were removed from the rostral sylvian mask. The raincloud plots[5] show the group mean task-based functional connectivity with V1 (black dots) measured in arbitrary units (a.u.) with error bars indicating the standard error of the mean (s.e.m.), individual means (coloured dots) and density plots (half violins). Planned comparisons were false discovery rate (FDR) corrected to control for multiple comparisons. *** $p < .001$; a.u. arbitrary units; s.e.m, standard error of the mean.

#### 2 Supplementary notes

##### Supplementary Note 1

##### Functional localizer: univariate results

Action observation led to activation in multiple temporal lobe regions in the dog brain (see e.g., **Figure 2A** for an overview of the dog action observation network). Due to mixed findings regarding face- and body-sensitive areas in the dog brain in past research, we used a functional localizer task to detect them in our sample. This allowed us to identify the functional analogue of the human ventral visual pathway [6] within the dog action observation network.

The agent localizer revealed greater activation in the mid suprasylvian gyrus both in response to static faces and bodies compared to inanimate objects (each condition > scrambled controls) but no difference in activation in this area between faces and bodies (henceforth referred to as mid suprasylvian animate area; see summary in **Figure 2B**; and **Supplementary Figure S8 and Table S3** for detailed results). Bodies further resulted in greater activation in the caudal suprasylvian gyrus than inanimate objects and faces (henceforth referred to as mid suprasylvian body area). Faces compared to bodies did not result in any significant activation. Observing conspecific (i.e., dog) faces and bodies compared to human faces and bodies resulted in greater activation in dogs' early visual and extrastriate cortices (i.e., marginal and ectomarginal gyri) and the mid suprasylvian animate area. The reversed contrast did not reveal any significant activation.

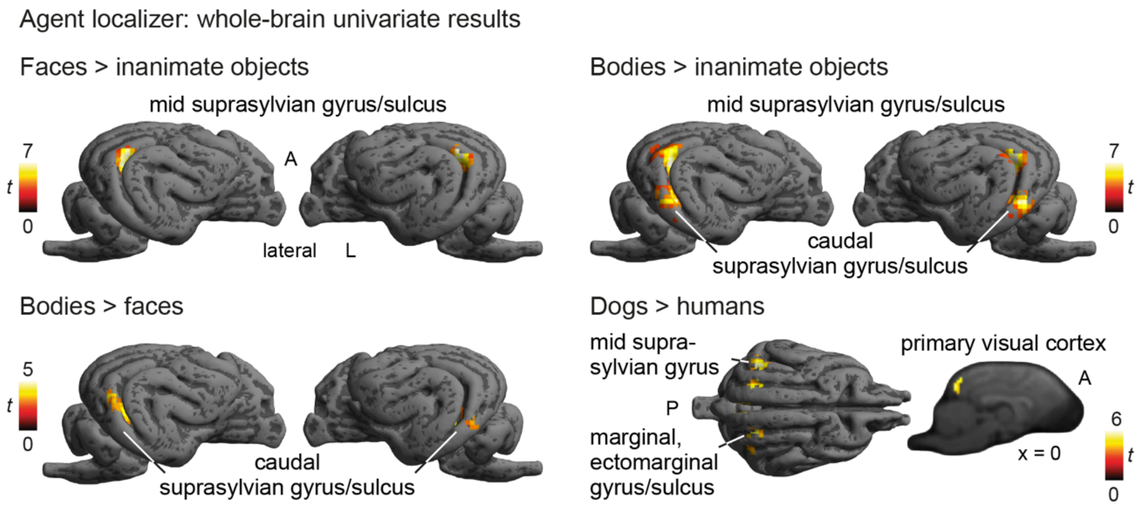


**Figure S8. Animate- and body-sensitive areas in the dog mid and caudal suprasylvian gyrus were identified using a functional localizer.** Results of the functional localizer indicate that static images of faces, as well as bodies compared to inanimate objects (all conditions > scrambled) resulted in greater activation in an overlapping area in the mid suprasylvian gyrus (upper row). Bodies compared to inanimate objects additionally revealed a significant bilateral cluster in the dog caudal suprasylvian gyrus, which also showed greater activation for bodies than faces (lower row, left). Observation of dog compared to human faces and bodies led to greater activation in early visual cortices (i.e., posterior marginal, splenial gyrus), the posterior ectomarginal gyrus and the face and body (i.e., agent-) sensitive mid suprasylvian area. The contrasts faces > bodies and humans > dogs did not reveal any significant clusters, and we did not find a significant interaction between agent (dog, human) and section (face, body). Results are $p < .05$ FWE-corrected at cluster-level using a cluster-defining threshold of $p < .005$; see also **Supplementary Table S3**. A, anterior; P, posterior; L, left; R, right; *t*, *t*-values.

#### Functional localizer: task-based functional connectivity with V1

To further investigate if the localized dog face- and body-sensitive areas (i.e., agent areas) are functional analogues of the human ventral visual pathway[6,7], we tested if the agent areas showed increased connectivity with V1, indicating increased exchange of information, during the perception of static faces and bodies compared to controls.

We found the strongest task-based functional connectivity between V1 and the temporal agent areas when dogs saw static images of faces and bodies compared to inanimate objects and scrambled controls (see **Supplementary Figure S9** and **Supplementary Table S11**). In the mid suprasylvian animate area, V1 connectivity did not differ between face and body perception, but in the caudal suprasylvian body area, functional connectivity was significantly stronger for bodies compared to faces. In the body area, connectivity during face perception was also significantly lower than inanimate object perception. Scrambled images resulted in the lowest V1 connectivity in both areas.

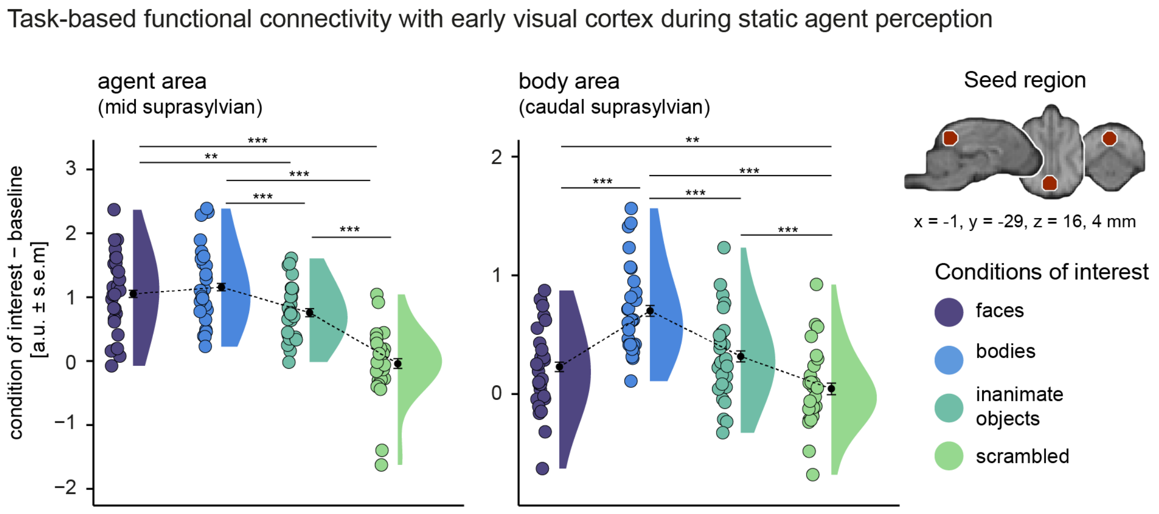


**Figure S9. Strongest task-based functional connectivity during agent perception compared to controls in agent and body area using localizer data.** Results show task-based functional connectivity with (V1) was significantly higher during face and body perception than control conditions. In the caudal suprasylvian body area, V1 connectivity was significantly higher during body compared to face perception. Connectivity measures between faces and bodies did not differ in the mid suprasylvian animate area. We also found significantly stronger V1 connectivity during inanimate object perception in both regions than the scrambled controls. In the body area, task-based connectivity during face perception was even significantly lower than during inanimate object perception (see **Supplementary Table S11**). The raincloud plots[5] show the group mean task-based functional connectivity with V1 (black dots) measured in arbitrary units (a.u.) with error bars indicating the standard error of the mean (s.e.m.), individual means (coloured dots) and density plots (half violins). Planned comparisons were false discovery rate (FDR) corrected to control for multiple comparisons. * $p < .05$, ** $p < .01$, *** $p < .001$; a.u., arbitrary units; s.e.m, standard error of the mean

### 3 Supplementary tables

**Table S1.** Dog action observation network

| Species, contrast & brain region | coordinates | | | *z*-value | cluster size |
|---|---|---|---|---|---|
| | x | y | z | | |
| **Action observation > implicit baseline** (*k* = 41) | | | | | |
| R mid suprasylvian gyrus | 17 | -24 | 14 | 6.46 | 2322 |
| **Action observation > object motion** (*k* = 39) | | | | | |
| L mid suprasylvian gyrus | -16 | -26 | 14 | 6.68 | 803 |
| R caudal suprasylvian gyrus | 18 | -24 | 2 | 5.51 | 135 |
| L rostral ectosylvian gyrus | -16 | -9 | 12 | 4.66 | 58 |
| R rostral ectosylvian sulcus | 20 | -4 | 8 | 4.41 | 71 |
| **Action observation > phase-scrambled actions** (*k* = 38) | | | | | |
| R rostral ectosylvian sulcus | 16 | -10 | 10 | 5.5 | 76 |
| L rostral ectosylvian sulcus | -16 | -10 | 12 | 5.1 | 169 |
| L mid suprasylvian gyrus | -16 | -24 | 12 | 4.63 | 72 |
| L caudal composite gyrus | -24 | -24 | -4 | 4.39 | 94 |
| R caudal suprasylvian gyrus | 18 | -24 | 1 | 4.15 | 66 |

*Note.* Effects were tested for significance with a cluster defining threshold of $p$ <.005 and a cluster probability threshold of $p$ < .05 FWE. We report the first local maximum within each cluster for each $t$-tests along with the critical cluster sizes ($k$). The data is presented in **Figure 2A-B and Supplementary Figure S3**. L, left; R, right.

**Table S2.** Human action observation network

| Contrast & brain region | coordinates | | | *z*-value | cluster size |
|---|---|---|---|---|---|
| | x | y | z | | |
| **Action observation > implicit baseline** (*k* = 59) | | | | | |
| R temporal occipital fusiform cortex | 42 | -52 | -18 | Inf | 15007 |
| R thalamus | 24 | -30 | 4 | Inf | 73 |
| R inferior frontal gyrus pars opercularis | 44 | 12 | 28 | 6.56 | 2674 |
| R amygdala | 30 | 0 | -18 | 6.28 | 193 |
| L precentral gyrus | -58 | 10 | 30 | 6.20 | 3513 |
| R superior parietal lobule | 32 | -48 | 56 | 6.18 | 1705 |
| L cerebellum | -10 | -74 | -22 | 5.47 | 66 |
| L frontal pole | -52 | 40 | -4 | 5.43 | 133 |
| R frontal pole | 50 | 40 | -12 | 4.58 | 129 |
| **Action observation > object motion** (*k* = 60) | | | | | |
| R occipital pole | 20 | -94 | 10 | Inf | 4351 |
| L inferior lateral occipital cortex | -46 | -80 | 4 | 7.56 | 3564 |
| R thalamus | 22 | -32 | 4 | 6.97 | 183 |
| R temporal pole | 46 | 8 | -26 | 4.84 | 167 |
| L frontal pole | 0 | 68 | 8 | 4.69 | 243 |
| L postcentral gyrus | -38 | -32 | 50 | 4.66 | 80 |
| L frontal medial cortex | -8 | 48 | -12 | 4.37 | 103 |
| R precuneus cortex | 6 | -56 | 28 | 4.3 | 64 |
| R postcentral gyrus | 36 | -38 | 62 | 4.21 | 181 |
| **Action observation > phase-scrambled actions** (*k* = 59) | | | | | |
| L inferior lateral occipital cortex | -40 | -86 | -8 | 7.73 | 3208 |
| R inferior lateral occipital cortex | 44 | -82 | -4 | 7.68 | 4160 |
| R intracalcarine cortex | 6 | -64 | 8 | 6.18 | 6785 |
| L cerebellum | -2 | -72 | -24 | 4.79 | 89 |
| R middle frontal gyrus | 38 | 4 | 60 | 4.77 | 3921 |
| L precentral gyrus | -56 | 4 | 40 | 4.73 | 105 |
| R amygdala | 28 | -2 | -18 | 4.52 | 62 |
| L precentral gyrus | -26 | -8 | 56 | 4.26 | 306 |
| R inferior frontal gyrus pars opercularis | 44 | 16 | 24 | 4.08 | 99 |

*Note.* Effects were tested for significance with a cluster defining threshold of $p < .001$ and a cluster probability threshold of $p < .05$ FWE. We report the first local maximum within each cluster for each *t*-tests along with the critical cluster sizes (*k*). The data is presented in **Figure 2A and Supplementary Figure S3**. L, left; R, right.

**Table S3.** Whole-brain results: agent localizer dogs

| Species, contrast & brain region | coordinates | | | *z*-value | cluster size |
|---|---|---|---|---|---|
| | x | y | z | | |
| **Faces > inanimate objects** (*k* = 41) | | | | | |
| L mid suprasylvian gyrus | -16 | -24 | 16 | 5.6 | 66 |
| R mid suprasylvian gyrus | 17 | -22 | 16 | 5.99 | 65 |
| **Bodies > inanimate objects** (*k* = 41) | | | | | |
| L caudal suprasylvian gyrus | -22 | -26 | 1 | 5.8 | 77 |
| R mid suprasylvian gyrus | 17 | -24 | 13 | 5.52 | 188 |
| L mid suprasylvian gyrus | -18 | -24 | 14 | 5.5 | 85 |
| **Bodies > faces** (*k* = 41) | | | | | |
| L caudal suprasylvian gyrus | -20 | -20 | 1 | 4.81 | 44 |
| R caudal suprasylvian gyrus | 18 | -24 | 4 | 4.29 | 71 |
| **Dogs > human** (*k* = 41) | | | | | |
| R mid suprasylvian gyrus | 16 | -26 | 18 | 4.32 | 94 |
| R splenial gyrus | 2 | -33 | 14 | 3.91 | 58 |
| L ectomarginal gyrus | -8 | -27 | 20 | 3.84 | 44 |

*Note.* Effects were tested for significance with a cluster defining threshold of $p$ <.005 and a cluster probability threshold of $p < .05$ FWE. We report the first local maximum within each cluster for each paired *t*-test along with the critical cluster sizes (*k*). The contrasts faces > bodies and humans > dogs did not reveal any significant clusters and we did not find a significant interaction between agent (dog, human) and section (face, body). The data is presented in **Figure 2C** and **Supplementary Figure S8**. L, left; R, right.

**Supplementary Table S4.** Univariate action observation activation dogs: LMM results[a]

| Anatomical region | F ($df_{\text{Num}}$, $df_{\text{Den}}$) | $p$ | $p_{\text{FDR}}$ | $\eta^2_g$ |
|---|---|---|---|---|
| Precruciate (premotor & SMA) | 1.74 (2,138) | .1786 | .3579 | .02 |
| Postcruciate (M1 & S1) | 1.31 (2,138) | .2739 | .3579 | .02 |
| Rostral suprasylvian (S1) | 4.56 (2,138) | **.012** | **.036** | .06 |
| Rostral ectosylvian (S2) | 18.46 (2,138) | **< .0001** | **.0003** | .21 |

*Note.* [a] Within-subjects design, dependent variable: activation levels, predictor: *condition* (levels: actions, object motion, phase-scrambled) and a random intercept for each subject. The model was estimated using restricted maximum likelihood. *P*-values for group comparisons are false discovery rate (FDR) corrected as they investigate the same research question in each region-of-interest (ROI); uncorrected *p*-values are also reported, and *p*-values < .05 are in bold. Post-hoc comparison results are presented in **Figure 4A**. SMA, supplementary motor area; M1, primary motor cortex; S1, primary somatosensory cortex; S2, secondary somatosensory cortex; $df_{Num}$, degrees of freedom numerator; $df_{Den}$ degrees of freedom denominator; $\eta^2_g$, generalized eta-squared.

**Supplementary Table S5.** Univariate action observation activation dogs: LMM results[a]

| Region-of-interest | F ($df_{Num}$, $df_{Den}$) | $p$ | $p_{FDR}$ | $\eta^2_g$ |
|---|---|---|---|---|
| Rostral suprasylvian (S1) | | | | |
| Action | .6 (1,81) | .4405 | .88 | 0 |
| Agent | .24 (1,81) | .6247 | .62 | 0 |
| Action × agent | .3 (1,81) | .5854 | .96 | 0 |
| Rostral ectosylvian (S2) | | | | |
| Action | 0 (1,81) | .92090 | .92 | 0 |
| Agent | 3.58 (1,81) | .06223 | .12 | 0 |
| Action × agent | 0 (1,81) | .96325 | .96 | 0 |

*Note.* [a]2 × 2 within-subjects design, dependent variable: activation levels, predictors: *agent* (levels: dog, human) and *action* (transitive, intransitive) with a random intercept for each subject. The model was estimated using restricted maximum likelihood. Analysis focused on regions-of-interest (ROIs) with greater activation for actions compared to controls (see **Supplementary Table S4**). *P*-values for group comparisons are false discovery rate (FDR) corrected as they investigate the same research question in each region-of-interest (ROI); uncorrected *p*-values are also reported, and *p*-values < .05 are in bold. S1, primary somatosensory cortex; S2, secondary somatosensory cortex; $df_{Num}$, degrees of freedom numerator; $df_{Den}$ degrees of freedom denominator; $\eta^2_g$, generalized eta-squared.

**Supplementary Table S6.** Cross-species comparison of lobe involvement: LMM results[a]

| Statistical map, predictor | F ($df_{Num}$, $df_{Den}$) | $p$ | $p_{FDR}$ | $\eta^2_g$ |
|---|---|---|---|---|
| **Cluster-corrected *t*-maps** (main analysis) | | | | |
| Lobe | 10.15 (1,66) | **.002** | **.003** | .13 |
| Sample | 24.68 (1,66) | **< .0001** | **< .0001** | .27 |
| Lobe × sample | 130.51 (1,66) | **< .0001** | **< .0001** | .66 |
| **Uncorrected *t*-maps** (secondary analysis) | | | | |
| Lobe | 4.16 (1,66) | **.045** | **.045** | .06 |
| Sample | 36.11 (1,66) | **< .0001** | **< .0001** | .35 |
| Lobe × sample | 92.07 (1,66) | **< .0001** | **< .0001** | .58 |
| **Top-5% voxels maps** (secondary analysis) | | | | |
| Lobe | 212.19 (1,66) | **< .0001** | **< .0001** | .76 |
| Sample | 87.56 (1,66) | **< .0001** | **< .0001** | .57 |
| Lobe × sample | 339.55 (1,66) | **< .0001** | **< .0001** | .84 |

*Note.* [a] 2 x 2 factorial design, dependent variable: percentage of active voxels, predictors: *lobe* (levels: parietal, temporal lobe), *sample* (levels: human, dog participants), and a random intercept for each subject. The model was estimated using restricted maximum likelihood. For the main analysis, significantly active voxels (i.e., statistical map) were determined via thresholding the individual contrast maps for the action observation contrast (i.e., all actions > implicit baseline) with a cluster-level threshold of $p$ < .005/.001 (dogs/humans) and a cluster probability of $p$ < .05 family-wise error (FWE). For the secondary control analyses, statistical maps were defined with a cluster-level threshold of $p$ < .005/.001 (dogs/humans) uncorrected or as the 5% most active voxels. *P*-values for group comparisons are false discovery rate (FDR) corrected as they investigate the same research questions for each statistical map; uncorrected *p*-values are also reported, and *p*-values < .05 are in bold. Post-hoc comparison results are presented in **Figure 4B** and **Supplementary Figure S6** fore secondary analyses**;** $df_{Num}$, degrees of freedom numerator; $df_{Den}$ degrees of freedom denominator; $\eta^2_g$, generalized eta-squared.

**Table S7.** 2 x 2 full factorial design: action (transitive, intransitive), agent (dog, human)

| Contrast & brain region | Coordinates | | | *z*-value | cluster size |
|---|---|---|---|---|---|
| | x | y | z | | |
| **Dog participants** | | | | | |
| **Conspecific (dog) > heterospecific (human) agent** | | | | | |
| L mid suprasylvian gyrus | -16 | -24 | 13 | 4.73 | 66 |
| **Human participants** | | | | | |
| **Transitive > intransitive action** | | | | | |
| L temporal occipital fusiform cortex | -30 | -56 | -16 | 7.96 | 880 |
| R temporal occipital fusiform cortex | 28 | -54 | -16 | 725 | 725 |
| R precentral gyrus | 22 | -14 | 58 | 4 | 81 |
| **Conspecific (human) > heterospecific (dog) agent** | | | | | |
| L occipital pole | -6 | -100 | 12 | Inf | 1461 |
| **Heterospecific > conspecific agent** | | | | | |
| L occipital fusiform gyrus | -22 | -90 | -10 | Inf | 3471 |
| R occipital pole | 18 | -92 | 2 | Inf | 4915 |
| L supramarginal gyrus, anterior division | -62 | -24 | 36 | 7.67 | 2554 |
| L precentral gyrus | -58 | 8 | 36 | 7.1 | 689 |
| R postcentral gyrus | 64 | -18 | 36 | 6.16 | 1783 |
| R superior parietal lobule | 34 | -48 | 62 | 4.65 | 665 |

*Note.* Effects were tested for significance with a cluster defining threshold of $p$ <.005/.001 (dogs/humans) and a cluster probability threshold of $p < .05$ FWE corrected for multiple (critical cluster size $k$ = 43/65 dogs/humans). We report the first local maximum within each cluster for all significant contrasts. The data is presented in **Figure 3**. L, left; R, right.

**Supplementary Table S8.** Lobe comparison task-based functional connectivity: LMM[a] results dogs

| Predictors | F ($df_{Num}$, $df_{Den}$) | *p* | $\eta^2_g$ |
|---|---|---|---|
| Condition | 6.75 (2, 30.36) | **.004** | .31 |
| Lobe | 83.34 (1, 27.06) | **< .0001** | .75 |
| Condition × lobe | 16.30 (2, 81) | **< .0001** | .29 |
| Contrasts for factor c*ondition* | t (*df*) | *p* | |
| actions – (object motion + scrambled motion) / 2 | 5.38 (81) | **< .0001** | / |
| scrambled motion – object motion | 1.92 (81) | .06 | / |

*Note.* [a] 2 × 3 within-subjects design, dependent variable: connectivity with primary visual cortex (V1) seed aggregated for each lobe, predictors: *condition* (levels: actions, object motion, scrambled motion) and *lobe* (temporal, parietal) with a random intercepts and slopes for *lobe* and *condition*. The factor condition was contrast coded using Helmert coding, the model was estimated using restricted maximum likelihood. Post-hoc comparison results are presented in **Figure 5A**. $df_{Num}$, degrees of freedom numerator; $df_{Den}$ degrees of freedom denominator; $\eta^2_g$, generalized eta-squared.

**Supplementary Table S9.** Lobe comparison task-based functional connectivity: LMM[a] results humans

| Predictors | F ($df_{Num}$, $df_{Den}$) | *p* | $\eta^2_g$ |
|---|---|---|---|
| Condition | 14.35 (2,39) | **< .0001** | .42 |
| Lobe | .13 (1,38) | .725 | 0 |
| Condition × lobe | 20.08 (2, 78) | **< .0001** | .34 |
| Contrasts for factor c*ondition* | t (*df*) | *p* | |
| actions – (object motion + scrambled motion) / 2 | 2.9 (78) | **.005** | / |
| scrambled motion – object motion | 5.63 (78) | **< .0001** | / |

*Note.* [a] 2 × 3 within-subjects design, dependent variable: connectivity with primary visual cortex (V1) seed aggregated for each lobe, predictors: *condition* (levels: actions, object motion, scrambled motion) and *lobe* (temporal, parietal) with a random intercepts and slopes for *lobe* and *condition*. The factor condition was contrast coded using Helmert coding, the model was estimated using restricted maximum likelihood. Post-hoc comparison results are presented in **Figure 5A**. $df_{Num}$, degrees of freedom numerator; $df_{Den}$ degrees of freedom denominator; $\eta^2_g$, generalized eta-squared.

**Supplementary Table S10**: Action observation task-based functional connectivity: LMM[a] results dogs

| Regions-of-interest | F ($df_{Num}$, $df_{Den}$) | $p$ | $p_{FDR}$ | $\eta^2_g$ |
|---|---|---|---|---|
| **Dog participants** | | | | |
| **Anatomically defined** | | | | |
| Mid suprasylvian gyrus (temporal) | 28.88 (2,54) | **< .0001** | **< .0001** | .52 |
| Caudal suprasylvian gyrus (temporal) | 14.47 (2,54) | **< .0001** | **< .0001** | .35 |
| Caudal composite gyrus (temporal) | 11.22 (2,54) | **< .0001** | **.0001** | .29 |
| Rostral sylvian gyrus (temporal) | 23.19 (2,54) | **< .0001** | **< .0001** | .46 |
| Anterior marginal gyrus (parietal) | .36 (2,54) | .7 | .73 | .01 |
| Anterior ectomarginal gyrus (parietal) | 6.82 (2, 54) | **.002** | **.003** | .2 |
| Presplenial gyrus (parietal) | .32 (2, 54) | .73 | .73 | .01 |
| **Functionally defined** (secondary analysis) | | | | |
| Mid suprasylvian animate area | 64.86 (2,54) | **< .0001** | **< .0001** | .71 |
| Caudal suprasylvian body area | 26.17 (2,54) | **< .0001** | **< .0001** | .49 |
| Caudal composite action area | 33 (2,54) | **< .0001** | **< .0001** | .55 |
| Rostral sylvian action area | (2,54) | **< .0001** | **< .0001** | .56 |
| **Human participants** | | | | |
| **Anatomically defined** | | | | |
| Fusiform cortex (temporal) | 16.46 (2,78) | **< .0001** | **< .0001** | .3 |
| Posterior superior temporal sulcus (temporal) | 22.63 (2,78) | **< .0001** | **< .0001** | .37 |
| Superior parietal lobule (parietal) | 17.12 (2,78) | **< .0001** | **< .0001** | .31 |
| Supramarginal gyrus (parietal) | 8.69 (2,78) | **.0004** | **.0004** | .18 |

*Note.* [a] Within-subjects design, dependent variable: task-based functional connectivity with primary visual cortex (V1) seed, predictor: *condition* (levels: actions, object motion, scrambled) with a random intercept for each subject. The model was estimated using restricted maximum likelihood. Regions-of-interest (ROIs) for the secondary analysis in the dog sample were defined based on the univariate results. All voxels overlapping with the ectosylvian gyrus (i.e., secondary somatosensory cortex) were removed from the rostral sylvian action area. *P*-values for group comparisons are false discovery rate (FDR) corrected as they investigate the same research question in each ROI; uncorrected *p*-values are also reported, and *p*-values < .05 are in bold. Post-hoc comparison results are presented in **Figure 5B and Supplementary Figure S7**. $df_{Num}$, degrees of freedom numerator; $df_{Den}$ degrees of freedom denominator; $\eta^2_g$, generalized eta-squared.

**Supplementary Table S11.** Agent localizer task-based functional connectivity: LMM results dogs

| Functionally defined region, predictor | F ($df_{Num}$, $df_{Den}$) | $p$ | $p_{FDR}$ | $\eta^2_g$ |
|---|---|---|---|---|
| (Mid suprasylvian) agent area[a] | | | | |
| Condition | 71.78 (3,81) | **< .0001** | **< .0001** | .73 |
| (Caudal suprasylvian) body area[a] | | | | |
| Condition | 37.64 (3, 81) | **< .0001** | **< .0001** | .58 |
| (Mid suprasylvian) agent area[b] | | | | |
| Agent | 12.05 (1,81) | **< .001** | **.002** | .13 |
| Section | 1.89 (1,81) | .17 | .17 | .02 |
| Section × agent | 1.13 (1,81) | .29 | .58 | .01 |
| (Caudal suprasylvian) body area [b] | | | | |
| Agent | 5.09 (1,81) | **.03** | **.03** | .06 |
| Section | 79.19 (1,81) | **< .0001** | **< .0001** | .49 |
| Section × agent | .31 (1,81) | .58 | .58 | .00 |

*Note.* [a]Within-subjects design, dependent variable: connectivity with primary visual cortex (V1) seed region, predictor: *condition* (levels: faces, bodies, inanimate objects, scrambled controls) with a random intercept for each subject. [b]2 × 2 within-subjects design, dependent variable: V1 connectivity, predictors: *agent* (levels: dog, human) and *section* (face, body) with a random intercept for each subject. Both models were estimated using restricted maximum likelihood. *P*-values for group comparisons are false discovery rate (FDR) corrected as they investigate the same research question in each region-of-interest (ROI); uncorrected *p*-values are also reported, and *p*-values < .05 are in bold. Post-hoc comparison results are presented in **Supplementary Figure S9**. $df_{Num}$, degrees of freedom numerator; $df_{Den}$ degrees of freedom denominator; $\eta^2_g$, generalized eta-squared.
